# The population genetics of human disease: the case of recessive, lethal mutations

**DOI:** 10.1101/091579

**Authors:** Carlos Eduardo G. Amorim, Ziyue Gao, Zachary Baker, José Francisco Diesel, Yuval B. Simons, Imran S. Haque, Joseph Pickrell, Molly Przeworski

**Author notes:** These authors co-supervised this work (Joseph Pickrell, Molly Przeworski). To whom correspondence should be addressed; Current address: Department of Ecology and Evolution, Stony Brook University, Stony Brook, NY.

## Abstract

Do the frequencies of disease mutations in human populations reflect a simple balance between mutation and purifying selection? What other factors shape the prevalence of disease mutations? To begin to answer these questions, we focused on one of the simplest cases: recessive mutations that alone cause lethal diseases or complete sterility. To this end, we generated a hand-curated set of 417 Mendelian mutations in 32 genes, reported to cause a recessive, lethal Mendelian disease. We then considered analytic models of mutation-selection balance in infinite and finite populations of constant sizes and simulations of purifying selection in a more realistic demographic setting, and tested how well these models fit allele frequencies estimated from 33,370 individuals of European ancestry. In doing so, we distinguished between CpG transitions, which occur at a substantially elevated rate, and three other mutation types. The observed frequency for CpG transitions is slightly higher than expectation but close, whereas the frequencies observed for the three other mutation types are an order of magnitude higher than expected. This discrepancy is even larger when subtle fitness effects in heterozygotes or lethal compound heterozygotes are taken into account. In principle, higher than expected frequencies of disease mutations could be due to widespread errors in reporting causal variants, compensation by other mutations, or balancing selection. It is unclear why these factors would have a greater impact on variants with lower mutation rates, however. We argue instead that the unexpectedly high frequency of disease mutations and the relationship to the mutation rate likely reflect an ascertainment bias: of all the mutations that cause recessive lethal diseases, those that by chance have reached higher frequencies are more likely to have been identified and thus to have been included in this study. Beyond the specific application, this study highlights the parameters likely to be important in shaping the frequencies of Mendelian disease alleles.

**Author Summary:** What determines the frequencies of disease mutations in human populations? To begin to answer this question, we focus on one of the simplest cases: mutations that cause completely recessive, lethal Mendelian diseases. We first review theory about what to expect from mutation and selection in a population of finite size and further generate predictions based on simulations using a realistic demographic scenario of human evolution. For a highly mutable type of mutations, such as transitions at CpG sites, we find that the predictions are close to the observed frequencies of recessive lethal disease mutations. For less mutable types, however, predictions substantially under-estimate the observed frequency. We discuss possible explanations for the discrepancy and point to a complication that, to our knowledge, is not widely appreciated: that there exists ascertainment bias in disease mutation discovery. Specifically, we suggest that alleles that have been identified to date are likely the ones that by chance have reached higher frequencies and are thus more likely to have been mapped. More generally, our study highlights the factors that influence the frequencies of Mendelian disease alleles.

## Introduction

New disease mutations arise in heterozygotes and either drift to higher frequencies or are rapidly purged from the population, depending on the strength of selection and the demographic history of the population [1–6]. Elucidating the relative contributions of mutation, natural selection and genetic drift will help to understand why disease alleles persist in humans. Answers to these questions are also of practical importance, in informing how genetic variation data can be used to identify additional disease mutations [7].

In this regard, rare, Mendelian diseases, which are caused by single highly penetrant and deleterious alleles, are perhaps most amenable to investigation. A simple model for the persistence of mutations that lead to Mendelian diseases is that their frequencies should reflect an equilibrium between their introduction by mutation and elimination by purifying selection, i.e., that they should be found at “mutation-selection balance” (MSB) [4]. In finite populations, random drift leads to stochastic changes in the frequency of any mutation, so demographic history, in addition to mutation and natural selection, also plays an important role in shaping the frequency distribution of deleterious mutations [3].

Another factor that may be important in determining the frequency of highly penetrant disease mutations is genetic interactions. For instance, a disease mutation may be rescued by another mutation in the same gene [8–10] or by a modifier locus elsewhere in the genome that modulates the severity of the disease symptoms or the penetrance of the disease allele (e.g. [11–13]).

For a subset of disease alleles that are recessive, an alternative model for their persistence in the population is that there is an advantage to carrying one copy but a disadvantage to carrying two or none, such that the alleles persist due to overdominance, a form of balancing selection. Well known examples include sickle cell anemia, thalassemia and G6PD deficiency in populations living where malaria is endemic [14]. The importance of overdominance in maintaining the high frequency of disease mutations is unknown beyond these specific cases.

Here, we tested hypotheses about the persistence of mutations that cause lethal, recessive, Mendelian disorders. This case provides a good starting point, because a large number of Mendelian disorders have been mapped (e.g., genes have already been associated with >66% of Mendelian disease phenotypes; [15]). Moreover, while the fitness effects of most diseases are hard to estimate, for recessive lethal diseases, the selection coefficient is clearly 1 for homozygote carriers in the absence of modern medical care (which, when available, became so only in the last couple of generations, a timescale that is much too short to substantially affect disease allele frequencies). Moreover, assuming mutation-selection balance in an infinite population would suggest that, given a per base pair (bp) mutation rate *u* on the order of 10^−8^ per generation [16], the frequency of such alleles would be 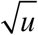, i.e., ~10^−4^ [4]. Thus, sample sizes in human genetics are now sufficiently large that we should be able to observe recessive disease alleles segregating in heterozygote carriers.

To this end, we compiled genetic information for a set of 417 mutations reported to cause fatal, recessive Mendelian diseases and estimated the frequencies of the disease-causing alleles from large exome datasets. We then compared these data to the expected frequencies of deleterious alleles based on models of MSB in order to evaluate the effects of demography and other mechanisms in influencing these frequencies.

## Results

### Mendelian recessive disease allele set

We relied on two datasets, one that describes 173 autosomal recessive diseases [17] and another from a genetic testing laboratory (Counsyl; <https://www.counsyl.com/>) that includes 110 recessive diseases of clinical interest. From these lists, we obtained a set of 44 “recessive lethal” diseases associated with 45 genes (Table S1), requiring that at least one of the following conditions is met: (i) in the absence of treatment, the affected individuals die of the disease before reproductive age, (ii) reproduction is completely impaired in patients of both sexes, (iii) the phenotype includes severe mental retardation that in practice precludes reproduction, or (iv) the phenotype includes severely compromised physical development, again precluding reproduction.

Based on clinical genetics datasets and the medical literature (see Methods for details), we were able to confirm that 417 Single Nucleotide Variants (SNVs) in 32 (of the 44) genes had been reported with compelling evidence of association to the severe form of the corresponding disease and an early-onset, as well as no indication of effects in heterozygote carriers (Table S2). By this approach, we obtained a set of mutations for which, at least in principle, there is no heterozygote effect, i.e., for which the dominance coefficient *h* = 0 in a model with relative fitness of 1 for the homozygote for the reference allele, 1-*hs* for the heterozygote, and 1-*s* for the homozygote for the deleterious allele, and the selective coefficient *s* is 1.

A large subset of these mutations (29.3%) consists of transitions at CpG sites (henceforth CpGti), which occur at a highly elevated rates (~17-fold higher on average) compared to other mutation types, namely CpG transversions, and non-CpG transitions and transversions [16]. This proportion is in agreement with previous estimates for a smaller set of disease genes [18] and for *DMD* [19].

### Empirical distribution of disease alleles in Europe

Allele frequency data for the 417 variants were obtained from the Exome Aggregation Consortium (ExAC) for 60,706 individuals, of whom 33,370 are non-Finnish Europeans [20]. Out of the 417 variants associated with putative recessive lethal diseases, three were found homozygous in at least one individual in this dataset (rs35269064, p.Arg108Leu in *ASS1*; rs28933375, p.Asn252Ser in *PRF1*; and rs113857788, p.Gln1352His in *CFTR*). Available data quality information for these variants does not suggest genotype calling artifacts (Table S2). Since these diseases have severe symptoms that lead to early death without treatment and these ExAC individuals are healthy (i.e., do not manifest severe Mendelian diseases) [20], the reported mutations are likely errors in pathogenicity classification or cases of incomplete penetrance (see a similar observation for *CFTR* and *DHCR7* in [21]). We therefore excluded them from our analyses. In addition to the mutations present in homozygotes, we also filtered out sites that had lower coverage in ExAC (see Methods), resulting in a final dataset of 385 variants in 32 genes (Table S2).

Genotypes for a subset (91) of these mutations were also available for a larger sample size (76,314 individuals with self-reported European ancestry) generated by the company Counsyl (Table S3). A comparison of the allele frequencies in this larger dataset to that of ExAC suggests that the allele frequencies for individual variants are concordant between the two datasets (Pearson's correlation coefficient of 0.79, Fig S1) and that the overall distributions do not differ appreciably (Kolmogorov–Smirnov test, p-value = 0.23). Thus, both data sets appear to reflect the general distribution of these disease alleles in Europeans. In what follows, we focus on ExAC, which includes a greater number of disease mutations.

### Models of mutation-selection balance

To generate expectations for the frequencies of these disease mutations under mutation-selection balance, we considered models of infinite and finite populations of constant size [3], and conducted forward simulations using a plausible demographic model for African and European populations [22] (see Methods for details). In all these models, there is a wild-type allele (A) and a deleterious allele (a, which could also represent a class of distinct deleterious alleles with the same fitness effect) at each site, such that the relative fitness of individuals of genotypes AA, Aa, or aa is given respectively by:

- *w_AA_*=1;
- *w_Aa_*=1-*hs*;
- *w_aa_*=1-*s*;

The mutation rate from A to a is *u*; we assume that there are no back mutations.

For a constant population of infinite size, Wright [23] showed that under these conditions, there exists a stable equilibrium between mutation and selection, when the selection pressure is sufficiently strong (*s*>>*u*). In particular, when the deleterious effect of allele a is completely recessive (*h*=0), its equilibrium frequency *q* is given by:

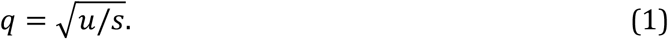

For a *finite* population of constant size, Nei [3] derived the mean (eq. 2) and variance (eq. 3) of the frequency of a fully recessive deleterious mutation (*h*=0) based on a diffusion model, leading to:

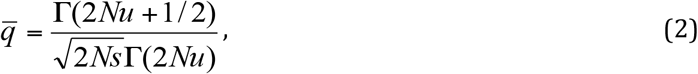

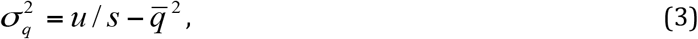

where *N* is the diploid population size and *Γ* is the gamma function (see Simons et al. [1] for a similar approximation).

In a finite population, the mean frequency, 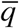, therefore depends on assumptions about the population mutation rate (2*Nu*). If the population mutation rate is high, such that 2*Nu*>>1, 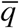 is approximated by

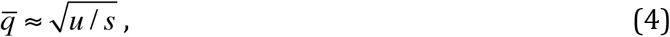

which is independent of the population size and equal to the equilibrium frequency in an infinite population, i.e., the right hand side of eq. (1). The important difference between the two models above is that in a finite population, there is a distribution of frequencies *q* (because of genetic drift), whose variance is given in eq. (3), rather than a single value, as in an infinite population.

In contrast, when the finite population has a low population mutation rate (*2Nu*<<1), the mean allele frequency, 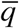, is approximated by:

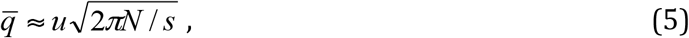

which depends on the population size [3].

We note that Nei [3] assumed a Wright-Fisher population, so there was no distinction between the census and the effective population size. However, when the two differ, it is the effective population size that governs the dynamics of deleterious alleles, so the *N* in the analytical results in fact represents the effective population size. In humans, the mutation rate at each bp is very small (on the order of 10^−8^ [16]) and the effective population size not that large, even recently [24,25], so the second approximation should apply when considering each single site independently.

The expectation and variance of the frequency of a fatal, fully recessive allele (i.e., *s*=1, *h*=0) are then given by:

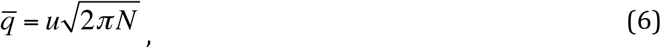

and

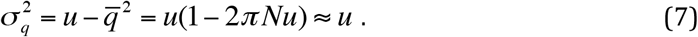

This single site model implicitly ignored the existence of compound heterozygosity in modeling the strength of selection acting on an individual site.

### Comparing mutation-selection balance models

Although an infinite population size has often been assumed when modeling deleterious allele frequencies (e.g. [5,26–29]), predictions under this assumption can differ markedly from what is expected from models of finite population sizes, assuming plausible parameter values for humans. For example, the long-term estimate of the effective population size from total polymorphism levels is ~20,000 individuals (assuming a mutation rate of 1.2 × 10^−8^ per bp per generation [16] and diversity levels of 0.1% [30]). In this case and considering a mutation rate of 1.5 × 10^−8^ for exons (which have a higher mutation rate than the rest of the genome, because of their base composition [31]), the average deleterious allele frequency in the model of finite population size is ~23-fold lower than that in the infinite population model (Fig 1).

**Fig 1.**
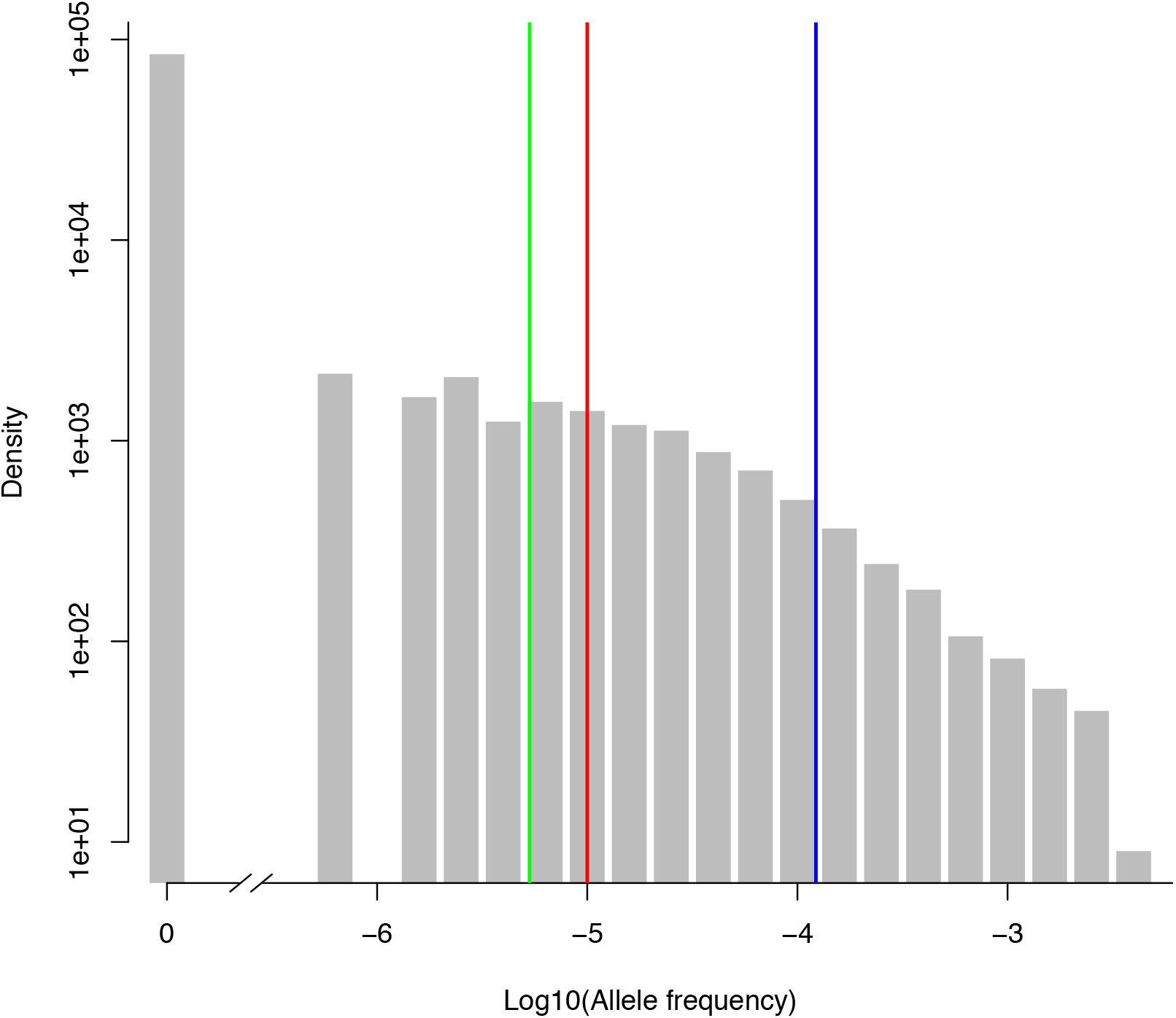
Expected allele frequencies in a population, according to mutation-selection balance models. The blue bar denotes the expected allele frequency under an infinite population size, the green bar the mean under a finite constant population, and the red bar the mean under a plausible demographic model for European populations (for this case, the entire distribution across 100,000 simulations is shown in the grey histogram). All models assume *s*=1 and *h*=0, i.e. fully recessive, lethal mutations. For the finite constant population size model, we present the mean frequency for a population size of 20,000 (see Fig S2A for other choices). Sample allele frequencies (*q*) were transformed to log_10_(*q*) and those *q*=0 were set to 10^−7^ for visual purposes, but indicated as “0” on the X-axis. The density of the distribution is plotted on a log-scale on the Y-axis. The mutation rate *u* was set to 1.5 × 10^−8^ per bp per generation for all models.

Because the human population size has not been constant and changes in the population size can affect the frequencies of deleterious alleles in the population (e.g. [2,32]), we also simulated the population dynamics of disease alleles under a plausible demographic model for European populations [22]. Assuming a mutation rate of 1.5 × 10^−8^ per bp, the mean allele frequency of a lethal, recessive disease allele obtained from this model was 9.91 × 10^−6^, ~1.86-fold higher than expected for a constant population size model with *N* = 20,000 (Fig 1). The mean frequency seen in simulations instead matches the expectation for a constant population size of 69,537 individuals (see Methods and Fig S2A). This finding is expected: the estimated effective population size of 20,000 is based on total genetic diversity; assuming that most of this variation is neutral, it therefore reflects an average timescale over millions of years. For recessive lethal mutations, which are relatively rapidly purged by natural selection, a more recent time depth is relevant (e.g., [1]). Indeed, in our simulations, most of the disease alleles (65.6%) segregating at present arose very recently, such that they were not segregating in the population 205 generations ago, the time point after which *Ne* is estimated to have increased from 9,300 to 512,000 individuals [22].

Increasing the effective population size in a constant size model is not enough to capture the dynamics of disease alleles appropriately, however. For example, if we compare simulation results obtained under the more complex Tennessen et al. [22] demographic model to those for simulations of a constant population size of *N* = 69,537, the mean allele frequencies match, but the distributions of allele frequencies are significantly different (Kolmogorov-Smirnov test, p-value < 10^−15^; Fig S2B and S2C). These findings thus confirm the importance of incorporating demographic history into models for understanding the population dynamics of disease alleles [5,33,34]. In what follows, we therefore test the fit of the more realistic demographic model [22] (and variants of it) to the observed allele frequencies.

### Comparing empirical and expected distributions of disease alleles

The mutation rate from wild-type allele to disease allele, *u*, is a critical parameter in predicting the frequencies of a deleterious allele [4, 35]. To model disease alleles, we considered four mutation types separately, with the goal of capturing most of the fine-scale heterogeneity in mutation rates [24,36–38]: transitions in methylated CpG sites (CpGti) and three less mutable types, namely transversions in CpG sites (CpGtv) and transitions and transversions outside a CpG site (nonCpGti and nonCpGtv, respectively). In order to control for the methylation status of CpG sites, we excluded 12 CpGti that occurred in CpG islands, which tend not to be methylated and thus are likely to have a lower mutation rate [36] (following Moorjani et al. [39]). To allow for heterogeneity in mutation rates within each one of these four classes considered, we modeled the within-class variation in mutation rates according to a lognormal distribution (see details in Methods and [24]).

For each mutation type, we then compared the mean allele frequency obtained from simulations to what is observed in ExAC, running 100,000 replicates. To this end, we matched simulations to the empirical data with regard to the number of individuals sampled and number of mutations observed of each mutation type and focused the analysis on the largest sample of the same common ancestry, namely Non-Finnish Europeans (*n* = 33,370) (Fig 2A). We find significant differences between empirical and expected mean frequencies for nonCpGti (50-fold higher on average; two-tailed p-value < 1.4 × 10^−4^; see Methods for details) and nonCpGtv (24-fold higher on average, two-tailed p-value < 1.2 × 10^−4^), to a lesser extent for CpGtv (9-fold higher on average, two-tailed p-value = 0.04). The mean frequency for CpGti is also somewhat higher than expected, but insignificantly so (2-fold higher on average, two-tailed p-value = 0.18). Intriguingly, the discrepancy between observed and expected frequencies becomes smaller as the mutation rate increases (Fig 2B).

**Fig 2.**
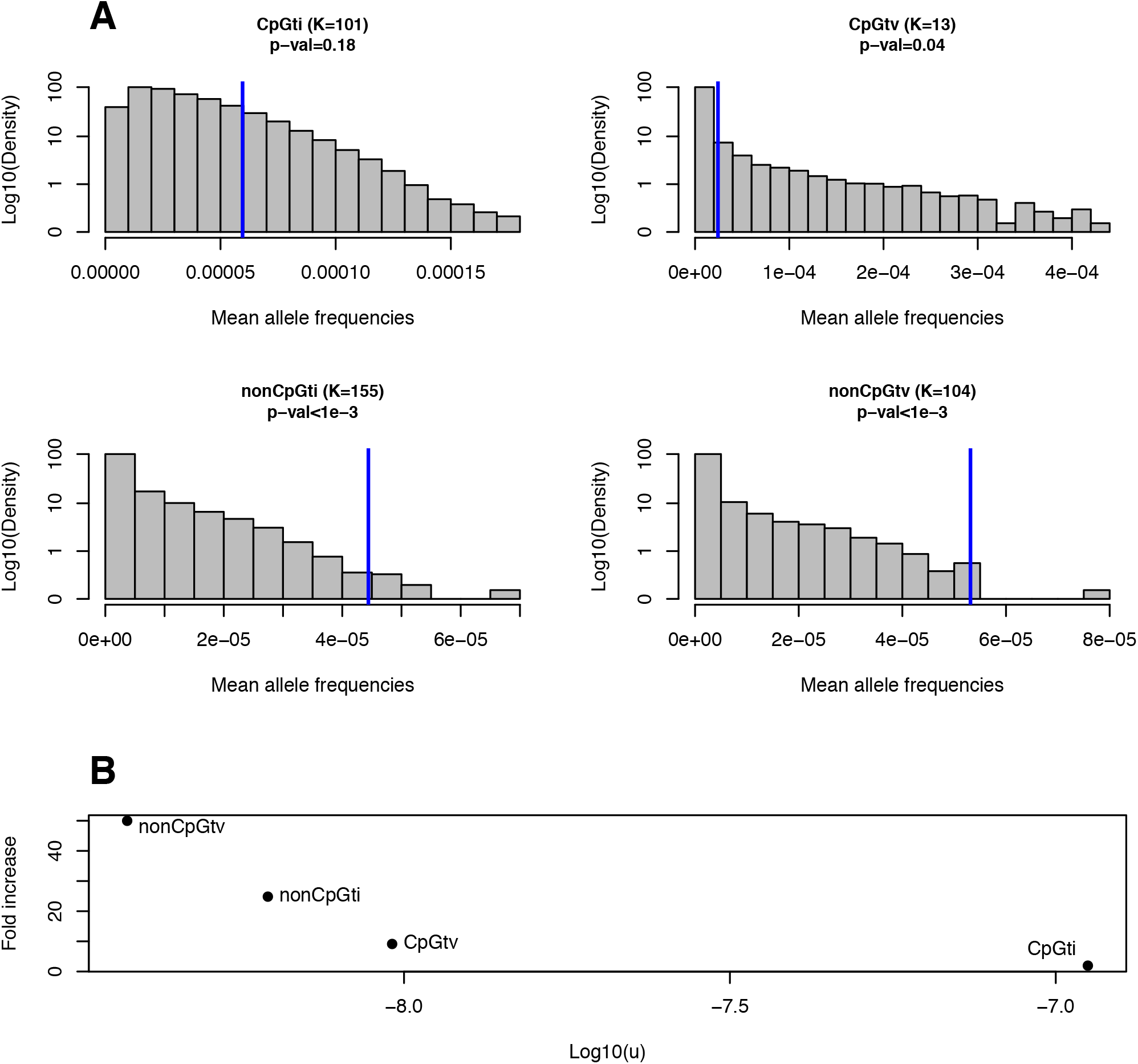
Comparison between expected and observed mean allele frequencies of recessive, lethal disease mutations. (A) Shown are the expected and observed frequencies of disease mutations for four different mutation types. The title of the panel indicates the mutation type, followed by *K*, the total number of mutations considered in this study of that type, with the p-value for the difference between observed and expected mean frequencies given below. Distributions in grey are the mean allele frequencies across *K* mutations based on 100,000 simulations, and rely on a plausible demographic model for European populations [22] (see Methods). Blue bars represent the observed mean frequencies of the four mutation types, estimated from 33,370 individuals of European ancestry from ExAC. (B) Fold increase in the observed mean allele frequency in relation to the expected, as a function of the mutation rate *u* (on a log-scale), for each of the four mutation classes.

Two additional factors that we have not included in our model should further decrease the predicted frequencies of disease alleles. Given that frequencies in ExAC are already unexpectedly high, these factors would only exacerbate the discrepancy between observed and expected frequencies of deleterious alleles. First, we have ignored the effects of compound heterozygosity, the case in which combinations of two distinct pathogenic alleles in the same gene lead to lethality. This phenomenon is known to be common [40], and indeed 58.4% of the 417 disease mutations considered in this study were initially identified in compound heterozygotes. In the presence of compound heterozygosity, each deleterious mutation will be selected against not only when present in two copies within the same individual, but also in the presence of lethal mutations at other sites of the same gene. Since the purging effect of selection against compound heterozygotes was not modeled in simulations, we would predict the frequency of a deleterious mutation to be even lower than shown (e.g., in Fig 2A).

In order to model the effect of compound heterozygosity in our simulations, we re-ran our simulations, but focusing on a gene rather than a single site and so considering the sum of frequencies of all known recessive lethal alleles within a gene. In these simulations, we used the same set-up as in the site level analysis, except for the mutation rate, *U*, which is now the sum of the mutation rates *u_j_* at each site *j* that is known to cause a severe and early onset form of the disease (Table S2; see Methods for details). This approach does not consider the contribution of other mutations in the genes that cause the mild and/or late onset forms of the disease, and implicitly assumes that all combinations of known recessive lethal alleles of the same gene have the same fitness effect as homozygotes. Comparing observed frequencies of disease alleles for each gene to predictions generated by simulation, one third of the 27 genes differ from the expected distribution at the 5% level, with a clear overall trend for observed frequencies to be above expectation (Table S4; Fig 3; Fisher's combined probability test p-value < 10^−14^).

**Fig 3.**
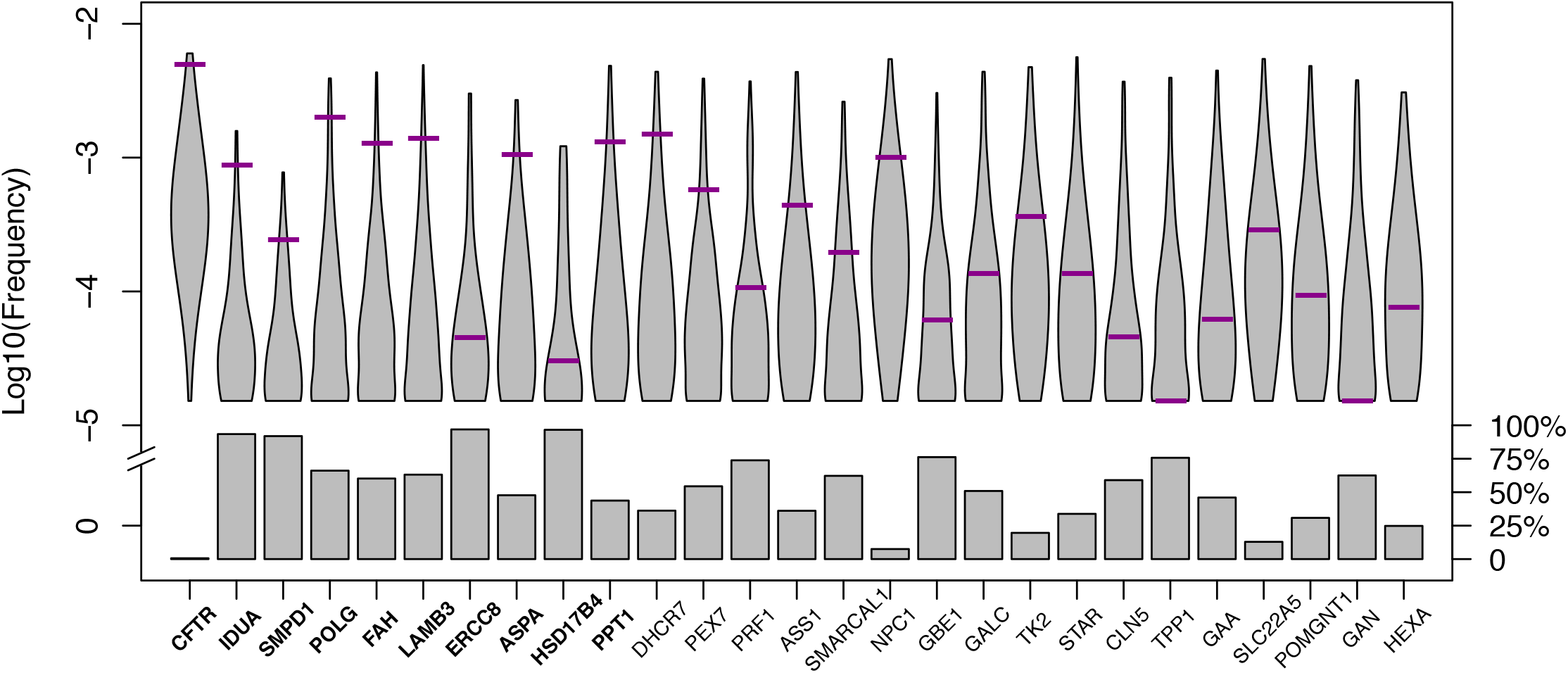
Disease allele frequencies at the gene level. The expectation (grey) is based on 1000 simulations, assuming no fitness decrease in heterozygotes, but allowing for compound heterozygosity (see Methods for details). The sum of allele frequencies of recessive lethal disease mutations in each gene (purple bars) was obtained from ExAC considering 33,370 European individuals. Genes are ordered according to the two-tailed p-value (Table S4; see Methods). Genes are bolded when they differ significantly from expectation (at the 5% level). Violin plots show the distribution of the log_10_ allele frequencies among segregating alleles obtained from simulations and boxes represent the fraction of simulations in which no deleterious allele was observed in the simulated sample at present.

This finding is even more surprising than it may seem, because we are far from knowing the complete mutation target for each gene, i.e., all the sites at which mutations could cause the disease. If there are additional, undiscovered sites in the gene at which mutations are fatal when carried in combination with a known recessive lethal mutation, the purging effect of purifying selection on the known mutations will be under-estimated in our simulations leading us to over-estimate the expected frequencies of the known mutations in simulations. Therefore, our predictions are, if anything, an over-estimate of the expected allele frequency, and the discrepancy between predicted and the observed is likely even larger than what we found.

The other factor that we did not consider in simulations but would reduce the expected allele frequencies is a subtle fitness decrease in heterozygotes. To evaluate potential fitness effects in heterozygotes when none had been documented in humans, we considered the phenotypic consequences of orthologous gene knockouts in mouse. We were able to retrieve information on phenotypes for both homozygote and heterozygote mice for only eight out of the 32 genes, namely *ASS1, CFTR, DHCR7, NPC1, POLG, PRF1, SLC22A5*, and *SMPD1*. For all eight, homozygote knockout mice presented similar phenotypes as affected humans, and heterozygotes showed a milder but detectable phenotype (Table S5). The magnitude of the heterozygote effect of these mutations in humans is unclear, but the finding with knockout mice makes it plausible that there exists a very small fitness decrease in heterozygotes in humans as well, potentially not enough to have been recognized in clinical investigations but enough to have a marked impact on the allele frequencies of the disease mutations. Indeed, even if the fitness effect in heterozygotes were as small as *h* = 1%, a 61% decrease in the mean allele frequency of the disease allele is expected relative to the case with complete recessivity (*h* = 0) (Fig S3).

## Discussion

To investigate the population genetics of human disease, we focused on mutations that cause Mendelian, recessive disorders that lead to early death or completely impaired reproduction. We sought to understand to what extent the frequencies of these mutations fit the expectation based on a simple balance between the input of mutations and the purging by purifying selection, as well as how other mechanisms might affect these frequencies. Many studies implicitly or explicitly compare known disease allele frequencies to expectations from mutation-selection balance [5, 26–29]. In this study, we tested whether known recessive lethal disease alleles as a class fit these expectations, and found that, under a sensible demographic model for European population history with purifying selection only in homozygotes, the expectations fit the observed disease allele frequencies poorly: the mean empirical frequencies of disease alleles are substantially above expectation for all mutation types (although not significantly so for CpGti), and the fold increase in observed mean allele frequency in relation to the expectation decreases with increased mutation rate (Fig 2). Furthermore, including possible effects of compound heterozygosity and subtle fitness decrease in heterozygotes will only exacerbate the discrepancy.

In principle, higher than expected disease allele frequencies could be explained by at least six (non-mutually exclusive) factors: (i) widespread errors in reporting the causal variants; (ii) misspecification of the demographic model, (iii) misspecification of the mutation rate; (iv) reproductive compensation; (v) overdominance of disease alleles; and (vi) low penetrance of disease mutations. Because widespread mis-annotation of the causal variants in disease mutation databases has previously been reported [20,41,42], we tried to minimize the effect of such errors on our analyses by filtering out any case that lacked compelling evidence of association with a recessive lethal disease, reducing our initial set of 769 mutations to 385 in which we had greater confidence (see Methods for details).

We also explored the effects of mis-specifying recent demographic history or the mutation rate. Based on very large samples, it has been estimated that population growth in Europe could have been stronger than what we considered in our simulations [43, 44]. When we consider higher growth rates, such that the current effective population size is up to 20-fold larger than that of the original model, we observe an increase in the expected frequency of recessive disease alleles and a larger number of segregating sites (Fig S4, columns A-F). However, the impact of larger growth rate is insufficient to explain the observed discrepancy: the allele frequencies observed in ExAC are still on average an order of magnitude larger than expected based on a model with a 20-fold larger current effective population size than the one initially considered [22] (Fig S4). Similarly, population substructure within Europe would only increase the number of homozygotes relative to what was modeled in our simulations (through the Wahlund effect [45]) and expose more recessive alleles to selection, thus decreasing the expected allele frequencies and exacerbating the discrepancy that we report.

In turn, to explore the effects of error in the mutation rate, we considered a 50% higher mean mutation rate than what has been estimated for exons [31], beyond what seems plausible based on current estimates on human mutation rates [39]. Except for the mean mutation rate (now set to 2.25 × 10^−8^), all other parameters used for these simulations (i.e. the variance in mutation rate across simulations, the demographic model [22], absence of selective effect in heterozygotes, and selection coefficient) were kept the same as the ones used for generating Fig S4 column A. The observed mean frequency remains significantly above what we predict using this high mutation rate and qualitative conclusions are unchanged (Fig S4, column G).

Another factor to consider is that for disease phenotypes that are lethal very early on in life, there may be partial or complete reproductive compensation (e.g. [46]). This phenomenon would decrease the fitness effects of the recessive lethal mutations and could therefore lead to an increase in the allele frequency in data relative to what we predict for a selection coefficient of 1. There are no reasons, however, for this phenomenon to correlate with the mutation rate, as seen in Fig 2B.

The other two factors, overdominance and low penetrance, are likely explanations for a subset of cases. For instance, *CFTR*, the gene in which mutations lead to cystic fibrosis, is the furthest above expectation (p-value = 0.002; Fig 3). It was long noted that there is an unusually high frequency of the *CFTR* deletion ΔF508 in Europeans, which led to speculation that disease alleles of this gene may be subject to over-dominance ([47, 48], but see [49]). Regardless, it is known that disease mutations in this gene can complement one another [8,9] and that modifier loci in other genes also influence their penetrance [9, 12]. Consistent with variable penetrance, Chen et al. [21] identified three purportedly healthy individuals carrying two copies of disease mutations in this gene. Similarly, *DHCR7*, the gene associated with the Smith-Lemli-Opitz syndrome, is somewhat above expectation in our analysis (p-value = 0.052; Fig 3) and healthy individuals were found to be homozygous carriers of putatively lethal disease alleles in other studies [21]. These observations make it plausible that, in a subset of cases (particularly for *CFTR*), the high frequency of deleterious mutations associated with recessive, lethal diseases are due to genetic interactions that modify the penetrance of certain recessive disease mutations. It is hard to assess the importance of this phenomenon in driving the general pattern that we observe, but two factors argue against it being a sufficient explanation for what we find at the level of single sites. First, when we remove 130 mutations in *CFTR* and 12 in *DHCR7*, the two genes that were outliers at the gene-level (Fig 3; Table S4) and for which we have evidence for incomplete penetrance [21], the discrepancy between observed and expected allele frequencies is barely impacted (Fig S5). Moreover, there is no obvious reason why the degree of incomplete penetrance would vary systematically with the mutation rate of a site, as we observe (Fig 2B).

Instead, it seems plausible that there is an ascertainment bias in disease allele discovery and mutation identification [48,50,51]. Unlike mis-sense or protein-truncating variants, Mendelian disease mutations cannot be readily annotated based solely on DNA sequences, and their identification requires reliable diagnosis of affected individuals (usually in more than one pedigree) followed by rigorous mapping of the underlying gene/mutation. Therefore, those mutations that have been identified to date are likely the ones that are segregating at higher frequencies in the population. Indeed, the guidelines for the interpretation of the pathogenicity of sequence variants by the American College of Medical Genetics and Genomics [52] may bias the identification of disease alleles towards high frequency ones. Moreover, mutation-selection balance models predict that the frequency of a deleterious mutation should correlate with the mutation rate. Together, these considerations suggest that disease variants of a highly mutable class, such as CpGti, are more likely to have been mapped and that the mean frequency of mapped mutations should be slightly above but close to that of all disease mutations in that class. We would further predict that, in contrast, less mutable disease mutations are less likely to have been discovered to date, and that the mean frequency of the subset of mutations that have been identified will far exceed that of all mutations in that class.

To quantify these effects, we modeled the ascertainment of disease mutations both analytically and in simulations (see Methods). As expected, we find that for a given mutation type, the average allele frequency of mutations that have been identified is always higher than that of all existing mutations (Table 1). Furthermore, comparison across different mutation types reveals that a higher mutation rate increases the probability of disease mutations being ascertained (Table 1 and Fig S6) and decreases the magnitude of bias in estimated allele frequency relative to the mutation class as a whole (Table 1). In summary, among all the possible aforementioned explanations for the observed discrepancy between empirical and expected mean allele frequencies, the ascertainment bias hypothesis is the only one that also explains why it is more pronounced for less mutable mutation types (Fig 2B).

**Table 1.** For a given set of parameters *n_a_* (the sample size in a putative disease ascertainment study) and *F_a_* (the average inbreeding coefficient of the population in which the study is being performed), we report the probability of a recessive, lethal mutation being ascertained, as well as the fold increase in mean allele frequencies of the ascertained cases (*q_a_*) in relation to the mean frequencies of all mutations for each mutation type (*q_u_*), based on simulations. See Methods for details. For a similar result derived from analytical modeling, see Fig S8. Parameters for this step of the simulation range according to plausible scenarios for human populations with widespread inbreeding (e.g., *F_a_*=1/16 corresponds to offspring of first-cousin marriage). The last column in the bottom panel shows the fold increase of the mean allele frequency observed in ExAC in relation to simulations based on the Tennessen et al. [22] demographic model (see Methods). Mutation rates *u* per bp, per generation were obtained from a large human pedigree study [16].

| Parameter | Simulation settings |  |  |  |  |  |
| --- | --- | --- | --- | --- | --- | --- |
| $n_a$ | 10,000 | 10,000 | 100,000 | 100,000 | 500,000 | ExAC |
| $F_a$ | 1/16 | 1/8 | 1/8 | 1/6 | 1/8 | |
| Mutation type ($u$) | Probability of a mutation being ascertained | | | | | |
| **CpGti (1.12e-7)** | 0.0150 | 0.0247 | 0.0968 | 0.1125 | 0.1858 | - |
| **CpGtv (9.59e-9)** | 0.0014 | 0.0024 | 0.0104 | 0.0124 | 0.0235 | - |
| **nonCpGti (6.18e-9)** | 0.0009 | 0.0015 | 0.0067 | 0.0074 | 0.0146 | - |
| **nonCpGtv (3.76e-9)** | 0.0005 | 0.0010 | 0.0044 | 0.0050 | 0.0091 | - |
|  | Fold increase in allele frequency due to ascertainment bias |  |  |  |  |  |
| **CpGti (1.12e-7)** | 27.6 | 22.3 | 9.0 | 8.0 | 5.2 | 1.9 |
| **CpGtv (9.59e-9)** | 284.1 | 222.6 | 81.2 | 70.2 | 40.6 | 9.2 |
| **nonCpGti (6.18e-9)** | 455.7 | 335.6 | 126.1 | 114.8 | 64.7 | 24.4 |
| **nonCpGtv (3.76e-9)** | 716.5 | 577.4 | 192.9 | 174.1 | 103.6 | 49.7 |

One clear implication of this hypothesis is that there are numerous sites at which mutations cause recessive lethal diseases yet to be discovered, particular at non-CpG sites. More generally, this ascertainment bias complicates the interpretation of observed allele frequencies in terms of the selection pressures acting on disease alleles. Beyond this specific point, our study illustrates how the large sample sizes now made available to researchers in the context of projects like ExAC [20] can be used not only for direct discovery of disease variants, but also to test why disease alleles are segregating in the population and to understand at what frequencies we might expect to find them.

## Methods

### Disease allele set

In order to identify single nucleotide variants within the 42 genes associated with lethal, recessive Mendelian diseases (Table S1), we initially relied on the ClinVar dataset [53] (accessed on June 3^rd^, 2015). We filtered out any variant that is an indel or a more complex copy number variant or that is ever classified as benign or likely benign in ClinVar (whether or not it is also classified as pathogenic or likely pathogenic). By this approach, we obtained 769 SNVs described as pathogenic or likely pathogenic. For each one of these variants, we searched the literature for evidence that it is exclusively associated to the lethal and early onset form of the disease and was never reported as causing the mild and/or late-onset form of the disease. We considered effects in the absence of medical treatment, as we were interested in the selection pressures acting on the alleles over evolutionary time scales rather than in the last one or two generations, i.e., the period over which treatment became available for some of diseases considered. To evaluate the impact of treatment, we decreased *s* from 1 to 0 (i.e., we assumed a complete absence of selective effects due to treatment) in the last three generations and compared the mean allele frequencies across 100,000 simulations implemented with or without this readjustment in selection coefficient. Because of the stochastic nature of the simulations, we repeated this pairwise comparison 10 times in order to get a range of expected increase in allele frequencies. We observe only a minor increase in the mean allele frequency (5.5% at most) across the 10 replicates. This simulation procedure corresponds to a scenario in which there is an extremely effective treatment for all diseases for the past three generations, which is a vast overestimate of the effect and length of treatment for the disease set that we consider.

Variants with mention of incomplete penetrance (i.e. for which homozygotes were not always affected) or with known effects in heterozygote carriers were removed from the analysis. This process yielded 417 SNVs in 32 genes associated with distinct Mendelian recessive lethal disorders (Table S2). Although these mutations were purportedly associated with complete recessive diseases, we sought to examine whether there would be possible, unreported effects in heterozygous carriers. To this end, we used the Mouse Genome Database (MGD) [54] (accessed July 29^th^, 2015) and were able to retrieve information for both homozygote and heterozygote mice for eight out of the 32 genes (all of which with a homologue in mice) (Table S5).

In addition to the information provided by ClinVar for each one of these variants, we considered the immediate sequence context of each SNV, to tailor the mutation rate estimate accordingly [16]. For doing so, we used an in-house Python script and the human genome reference sequence hg19 from UCSC (<http://hgdownload.soe.ucsc.edu/goldenPath/hg19/chromosomes/>).

### Genetic datasets

The Exome Aggregation Consortium (ExAC) [20] was accessed on August 9^th^, 2016. The data consist of genotype frequencies for 60,706 individuals, assigned after Principal Component Analysis to one of seven population labels: African (n=5,203), East Asian (n=4,327), Finnish (n=3,307, Latino (n=5,789), Non-Finnish European (n=33,370), South Asian (n=8,256) and “other” (n=454) [20]. We focused our analyses on those individuals of Non-Finnish European descent, because they constitute the largest sample size from a single ancestry group. We note that, some diseases mutations, for instance, those in *ASPA*, *HEXA* and *SMPD1*, are known to be especially prevalent in Ashkenazi Jewish populations, which could potentially bias our results if Ashkenazi Jewish individuals constituted a great portion of the sample we considered. However, this sample includes only ~2,000 (~6%) Ashkenazi individuals (Dr. Daniel MacArthur, personal communication).

From the initial 417 mutations, we filtered out three that were homozygous in at least one individual in ExAC and 29 that had lower coverage, i.e., fewer than 80% of the individuals were sequenced to at least 15x. This approach left us with a set of 385 mutations with a minimum coverage of 27x per sample and an average coverage of 69x per sample (Table S2). For 248 sites with non-zero sample frequencies, ExAC reported the number of non-Finnish European individuals that were sequenced, which was on average 32,881 individuals [20]. For the remaining 137 sites, we did not have this information. Nonetheless, the mean coverage across all samples is reported for each site and does not differ between the two sets of sites (Fig S7). We therefore assumed that mean number of individuals covered for all sites was 32,881 [55] and used this number to obtain sample frequencies from simulations, as explained below.

A second genetic dataset was obtained from Counsyl (<https://www.counsyl.com/>). Counsyl is a commercial genetic screening laboratory that offers, among other products, the “Family Prep Screen”, a genetic screening test intended to detect carrier status for up to 110 recessive Mendelian diseases in couples that are planning to have a child. A subset of 294,000 of its customers was surveyed by genotyping or sequencing for “routine carrier screening”. This subset excludes individuals with indications for testing because of known personal or family history of Mendelian diseases, infertility, and consanguinity. It therefore represents a more random (with regard to the presence of disease alleles), population-based survey. For these individuals, we had details on self-reported ancestry (14 distinct ethnic/ancestry/geographic groups) and the allele frequencies for 98 mutations that match those that passed our variant selection criteria described above, of which 91 are also sequenced to high coverage in the ExAC database (Table S2). We focused our analysis of this dataset on the 76,314 individuals with self-reported Northern or Southern European ancestry.

### Simulating the evolution of disease alleles with population size change

We modeled the frequency of a deleterious allele in human populations by forward simulations based on a crude but plausible demographic model for human populations from Africa and Europe, inferred from exome data for African-Americans and European-Americans [22]. To this end, we used a program described in [1]. In brief, the demographic scenario consists of an Out-of-Africa demographic model, with changes in population size throughout the population history, including a severe bottleneck in Europeans following the split from the African population and a rapid, recent population growth in both populations [22]. As in Simons et al. [1], we simulated genetic drift and two-way gene flow between Africans and Europeans in recent history. Negative selection acting on a single bi-allelic site was modeled as in the analytic models.

Allele frequencies follow a Wright-Fisher sampling scheme in each generation according to these viabilities, with migration rate and population sizes varying according to the demographic scenario considered. Whenever a demographic event (e.g. growth) altered the number of individuals and the resulting number was not an integer, we rounded it to the nearest integer, as in Simons et al. [1]. A burn-in period of 10*Ne* generations with constant population size *Ne* = 7,310 individuals was implemented in order to ensure an equilibrium distribution of segregating alleles at the onset of demographic changes in Africa, 148 Kya.

In contrast to Simons et al. [1], our simulations always start with the ancestral allele A fixed and mutation occurs exclusively from this allele to the deleterious one (a), i.e. a mutation occurs with mean probability *u* per gamete, per generation, and there is no back-mutation. However, recurrent mutations at a site are allowed, as in Simons et al. [1].

When implementing the model, we considered a mean mutation rate *u* of 1.5×10^−8^ per bp, per generation, as has been estimated for exons [31], as well as mutation rates for four distinct mutation types (CpGti = 1.12 × 10^−7^; CpGtv = 9.59 × 10^−9^; nonCpGti = 6.18 × 10^−9^; and nonCpGtv = 3.76 × 10^−9^) estimated from a large human pedigree study [16]. While these four categories capture much of the variation in germline mutation rates across sites, a number of other factors (e.g., the larger sequence context or the replication timing) also influence mutation rates, introducing heterogeneity in the mutation rate within each class considered [24,36,37,56]. To allow for this heterogeneity as well as for uncertainty in the point mutation rates estimates, in each simulation, instead of using a fixed rate *u* for each mutation type, we drew the mutation rate *M* from a lognormal distribution with the following parameters:

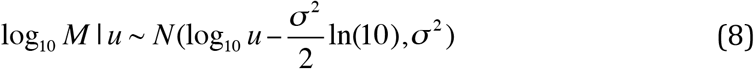

such that that *E*[*M*] = *u*. σ was set to 0.57 (following [24]).

For each mutation type, we then proceeded as follows:

1. We ran two million simulations, thus obtaining the distribution of deleterious allele frequencies expected for the European population.
2. We sampled *K* allele frequencies from the two million simulations implemented for each mutation type, where *K is* the number of identified mutations of that type. Sample allele frequencies were simulated from these population frequencies by Poisson sampling, so to match ExAC's number of chromosomes.
3. We repeated step (2) 100,000 times, thus obtaining a distribution for the mean allele frequency across *K* mutations.

To assess the significance of the deviation between observed and expected mean, we obtained a two-tailed p-value, defined as 2 × (*r*+1)/(100000+1), where *r* is the number of simulated allele frequencies that were greater or equal to that of the empirical mean [57], for each mutation type separately.

A well-known source of heterogeneity in mutation rate within the CpGti class is methylation status, with a high transition rate seen only at methylated CpGs [18]. In our analyses, we tried to control for the methylation status of CpG sites by excluding sites located in CpG islands (CGIs), which tend to not be methylated [39]. The CGI annotation for hg19 was obtained from UCSC Genome Browser (track “Unmasked CpG”; <http://hgdownload.soe.ucsc.edu/goldenPath/hg19/database/cpgIslandExtUnmasked.txt.gz>, accessed in June 6th, 2016). BEDTools [58] was used to exclude those CpG sites located in CGIs. We note that the CpGti estimate from [16] includes CGIs, and in that sense the average mutation rate that we are using for CpGti may be a very slight underestimate of the mean rate for transitions at methylated CpG sites.

Unless otherwise noted, the expectation assumes fully recessive, lethal alleles with complete penetrance. Notably, by calculating the expected frequency one site at a time, we are ignoring possible interaction between genes (i.e., effects of the genetic background) and among different mutations within a gene (i.e., compound heterozygotes). These assumptions are relaxed in two ways. In one analysis (Fig S3), we consider a very low selective effect in heterozygous individuals (*h* = 1%), reasoning that such an effect could plausibly go undetected in medical examinations and yet would nonetheless impact the frequency of the disease allele. Second, when considering the gene-level analysis (Fig 3), we implicitly allow for compound heterozygosity between any pair of known lethal mutations. For this analysis, we ran 1000 simulations for a total mutation rate *U* per gene that was calculated accounting for the heterogeneity and uncertainty in the mutation rates estimates as follows: (i) For *j* sites in a gene known to cause a recessive lethal disease and that passed our filtering criteria (Table S2), we drew a mutation rate *u_j_* from the lognormal distribution, as described above; (ii) We then took the sum of *u_j_* as the total mutation rate *U*; (iii) We then ran one replicate with *U* as the mutation parameter, and other parameters as specified for site level analysis. Because the mutational target size considered in simulations is only comprised of those sites at which mutations are known to cause a lethal recessive disease, it is almost certainly an underestimate of the true mutation rate—potentially by a lot. We note further that by this approach, we are assuming that compound heterozygotes formed by any two lethal alleles have fitness zero, i.e., that they are identical in their effects to homozygotes for any of the lethal alleles. Moreover, we are implicitly ignoring the possibility of complementation, which is (somewhat) justified by our focus on mutations with severe effects and complete penetrance (but see Discussion). Since we were interested in understanding the effect of compound heterozygosity, for this analysis, we did not consider the five genes in which only one mutation passed our filters (*BCS1L, FKTN, LAMA3, PLA3G6*, and *TCIRG1*).

### Modeling the effect of the ascertainment bias in disease discovery

To calculate the probability of ascertaining a recessive, lethal mutation, we assumed that all currently known disease mutations were identified in a putative ascertainment study of sample size *n_a_* in a population with an inbreeding coefficient of *F_a_*. Under this model, we can estimate *P_a_*, the probability of ascertaining a disease mutation, as following:

For a disease allele (denoted as *a*) at frequency *p* in the present population, if we randomly sample an individual with inbreeding coefficient of *F*, the probabilities of the three genotypes are:

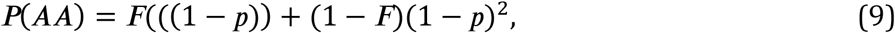

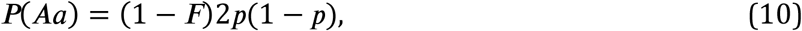

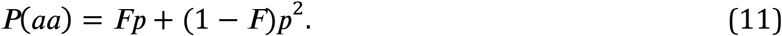

Thus, if *n_a_* unrelated individual are surveyed, the probability of not seeing any homozygote for the deleterious allele (which is the same as the probability of not being ascertained in this set) is:

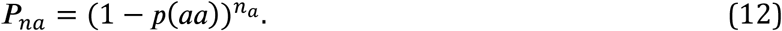

Therefore, the probability of ascertainment is

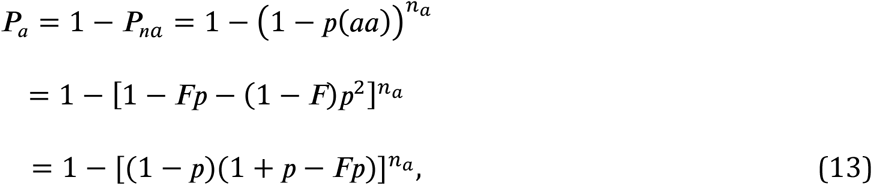

which is an increasing function with regard to *p* (the population allele frequency), *F* (the inbreeding coefficient of the population under study) as well as *n_a_* (the sample size of the putative ascertainment study) (Fig S6).

We also demonstrate the relationship between the probability of ascertainment and mutation rate using simulations of ascertainment bias implemented according to the following steps:

1. For each of the four mutation types considered, we generated 10^6^ allele frequencies *q* from the results of the simulations based on a realistic demographic model [22].
2. We generated *n_a_* independent diploid genotypes, given the allele frequencies from step 1 and an inbreeding coefficient *F_a_*. We ran this step for a range of *n_a_* and *F_a_* values (Table 1).
3. With a given combination of *n_a_* and *F_a_* values, we identified the cases (out of the 10^6^ observations from step 1) where at least one homozygote individual was observed in step 2. These cases correspond to disease mutations that were ascertained; the reasoning being that given complete penetrance, a recessive disease mutation can only be identified if there is at least one affected individual in the studied population. With this step, we calculated the probability of ascertainment by taking the fraction of cases that satisfy the criteria above.
4. Finally, for each one of the 10^6^ simulations from step 1, we generated a sample allele frequency of the disease mutation, matching ExAC's sample size (i.e., considering 2n = 65,762 chromosomes). We can then compare *q_u_*, the unbiased average allele frequency of all disease mutations, to *q_a_*, the mean frequency of the subset of cases ascertained in step 3, i.e., those cases for which at least one homozygote individual is observed.

These simulations were meant to illustrate the likely impact of ascertainment bias, rather than to precisely describe the disease mutation identification process or to quantify the expected effect. Notably, we performed these simulations for single sites, so the criteria for ascertainment in step 3 did not include the possibility of compound heterozygotes, despite the fact that most (58.4%) of the 417 disease mutations included in our study were initially identified in compound heterozygotes. However, this simulation framework could readily be extended in this direction and it would not change our qualitative conclusion.

## Acknowledgements

We thank Daniel MacArthur for his help with the ExAC data, Ellen Leffler for providing her Python script (available at https://github.com/cegamorim/PopGenHumDisease), as well as Aravinda Chakravarti, Damien Labuda and four anonymous reviewers for comments on an earlier version of the manuscript. All codes and data to generate the figures in R [59] and the script used to get the sequence context of each mutation are available at https://github.com/cegamorim/PopGenHumDisease. The code to run the simulations is available at https://github.com/sellalab/ForwardSimulator. Allele frequencies and other information for the disease mutations employed in the analyses are in Tables S2 and S3.

## Author Contributions

Conceived the study: JP, MP. Designed the study: CEGA, ZG, MP. Analyzed the data: CEGA. Implemented analytical models: ZG. Wrote the paper: CEGA, ZG, MP. Helped in acquisition and analysis of data: ZB, JFD. Contributed analytical tools or data: YBS, IR.

## Supporting information

**Table S1.**
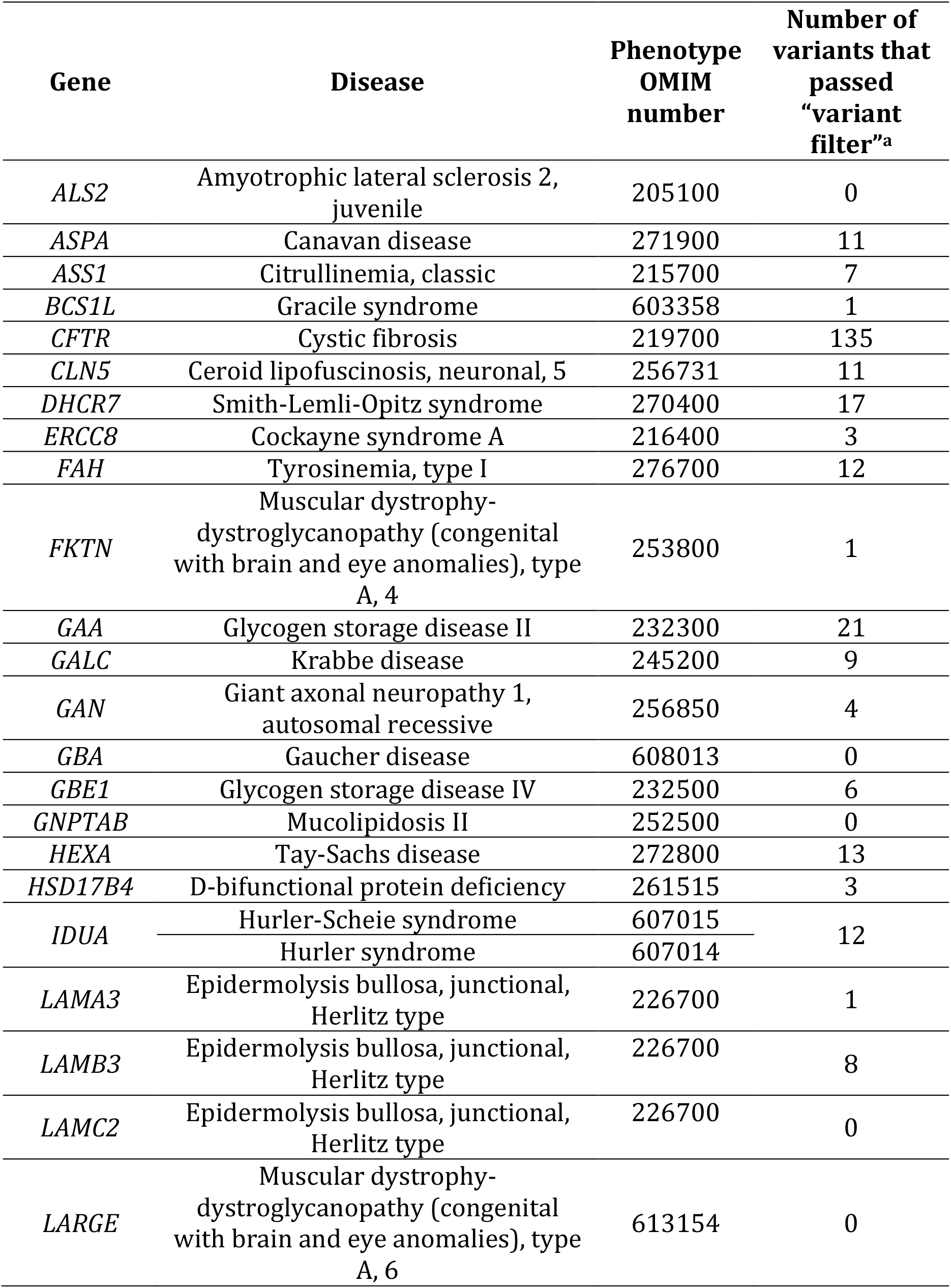
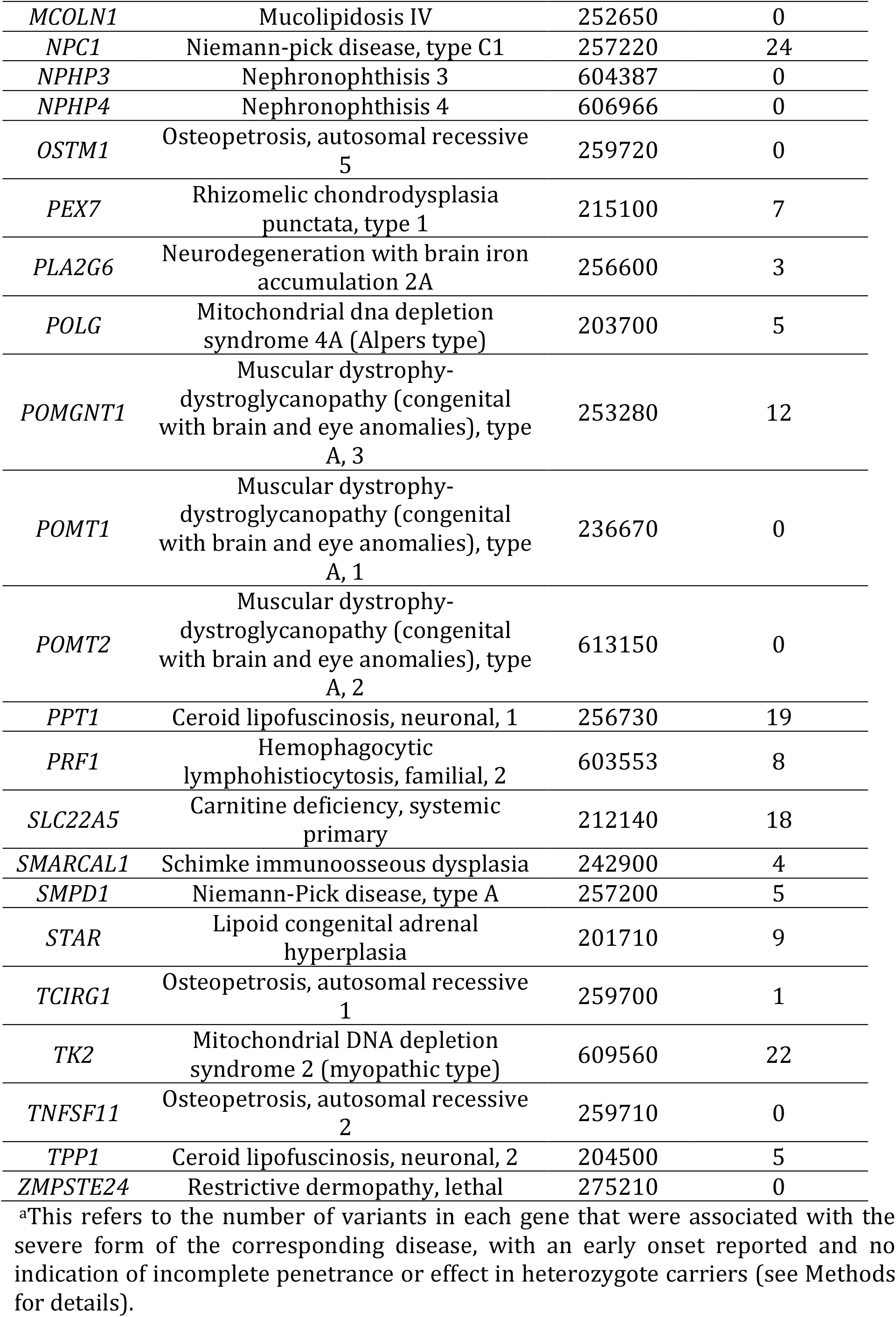
List of lethal, recessive Mendelian diseases.

**Table S2. Information on 417 mutations associated with the severe form of lethal, recessive Mendelian diseases.**

*Available at https://github.com/cegamorim/PopGenHumDisease*

**Table S3. Information on 91 mutations associated with the severe form of lethal, recessive Mendelian diseases in Counsyl and ExAC databases.**

*Available at https://github.com/cegamorim/PopGenHumDisease*

**Table S4.** P-values for the deviation of expected frequencies of deleterious alleles per gene, in comparison to the expected under SIM.

| **Gene** | **P-value** |
| --- | --- |
| *ASPA* | 0.032 |
| *ASS1* | 0.144 |
| *CFTR* | 0.002 |
| *CLN5* | 0.456 |
| *DHCR7* | 0.052 |
| *ERCC8* | 0.03 |
| *FAH* | 0.02 |
| *GAA* | 0.476 |
| *GALC* | 0.3 |
| *GAN* | 0.75 |
| *GBE1* | 0.218 |
| *HEXA* | 0.85 |
| *HSD17B4* | 0.044 |
| *IDUA* | 0.002 |
| *LAMB3* | 0.02 |
| *NPC1* | 0.194 |
| *PEX7* | 0.054 |
| *POLG* | 0.018 |
| *POMGNT1* | 0.62 |
| *PPT1* | 0.048 |
| *PRF1* | 0.132 |
| *SLC22A5* | 0.496 |
| *SMARCAL1* | 0.144 |
| *SMPD1* | 0.01 |
| *STAR* | 0.454 |
| *TK2* | 0.318 |
| *TPP1* | 0.464 |

**Table S5.**
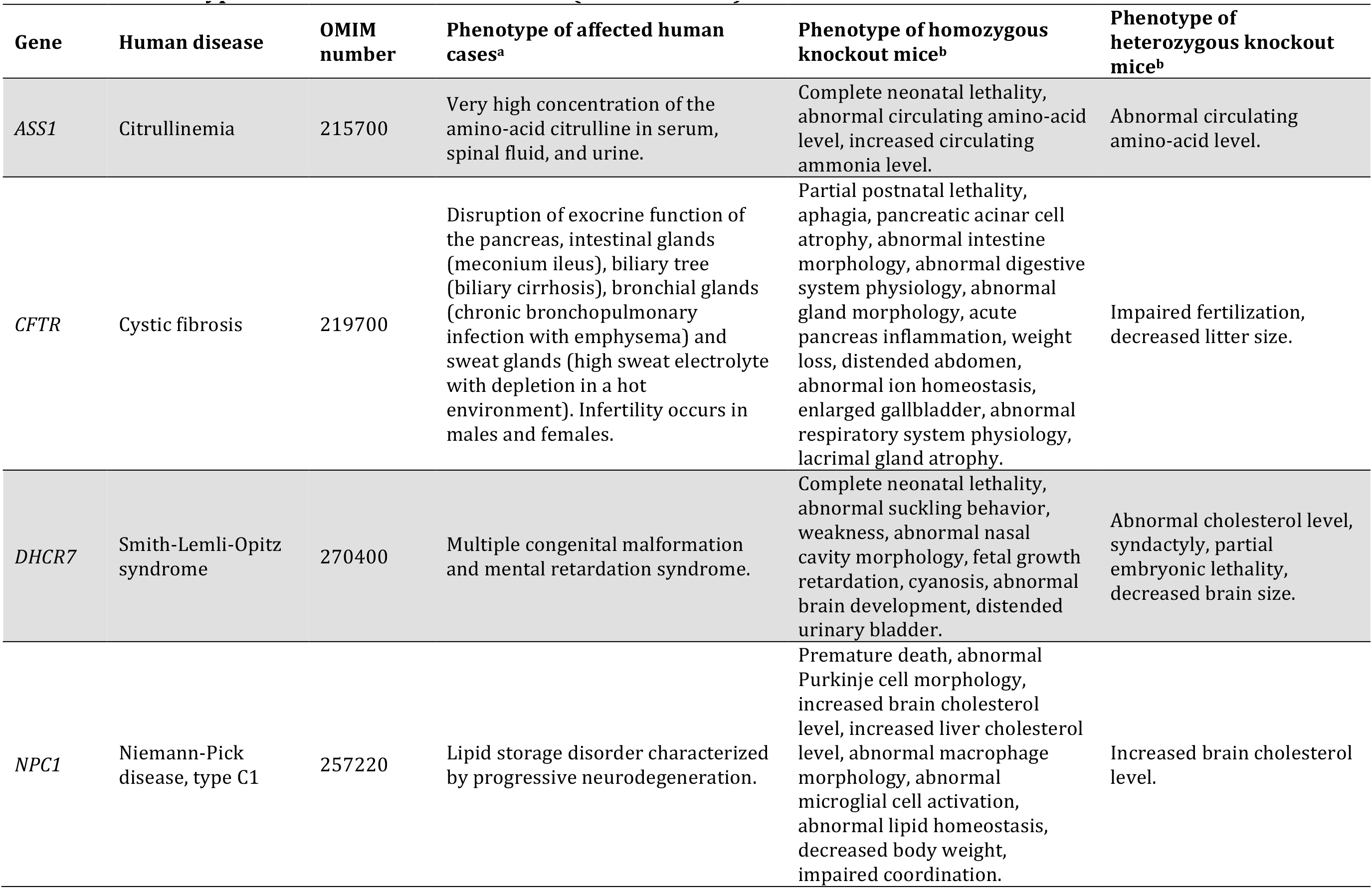
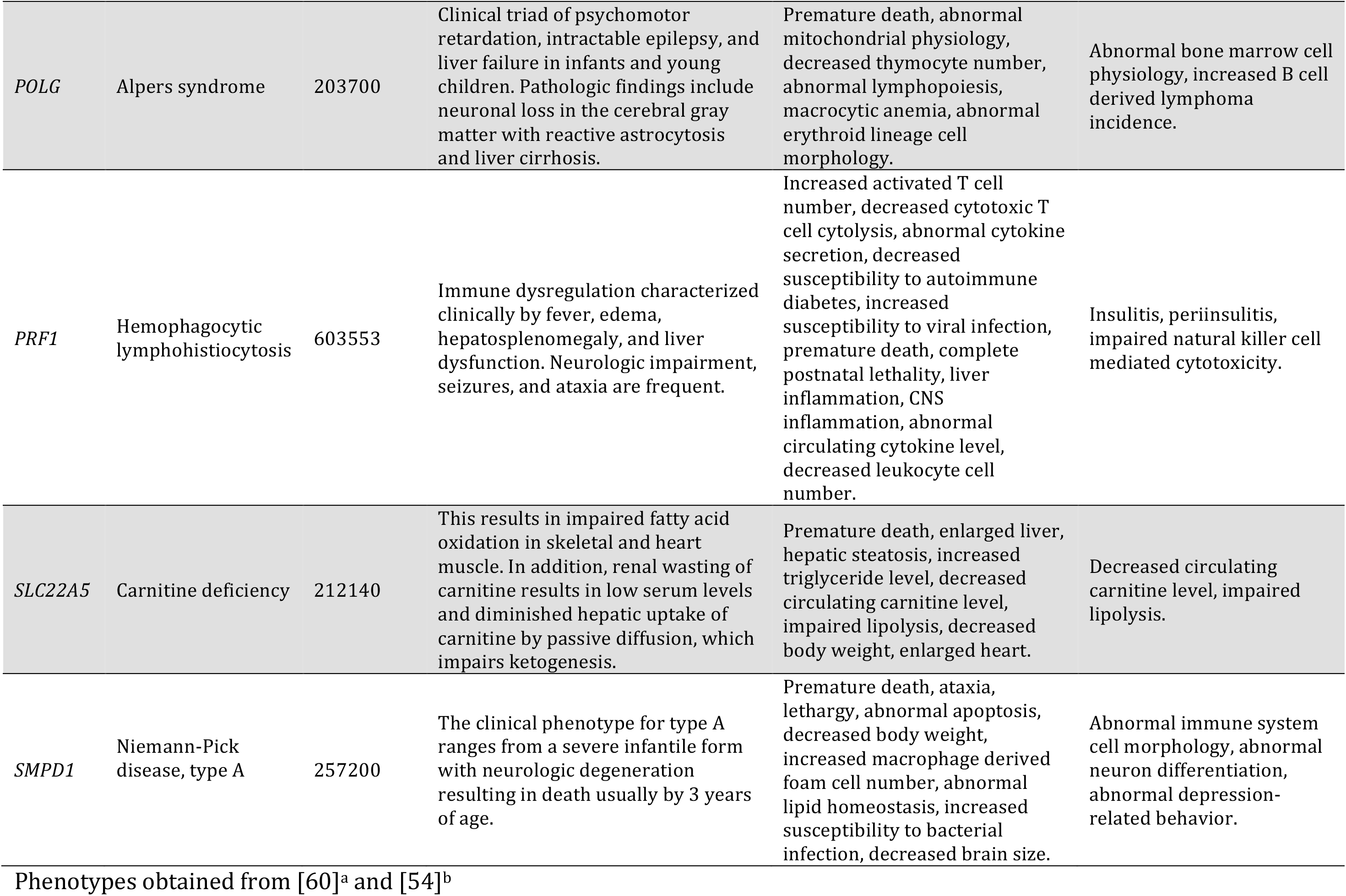
Phenotypic effect of mouse knock-outs (see main text)

**Fig S1.**
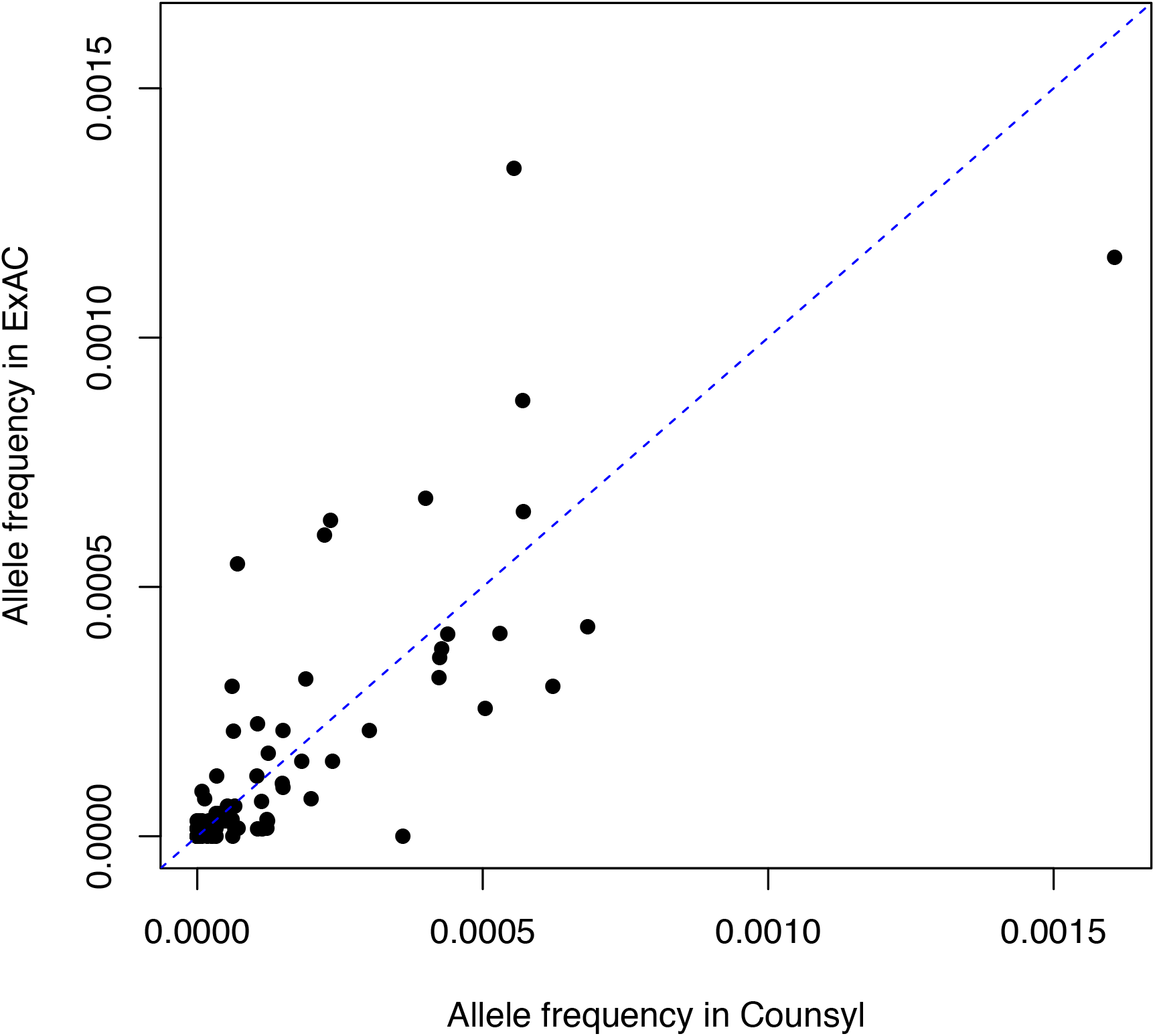
Comparison of the empirical allele frequency of recessive, lethal disease mutations in individuals of European ancestry from two large exome studies. Shown are the allele frequencies for 91 variants associated with lethal, recessive diseases, as estimated from 33,370 individuals of non-Finnish, European ancestry in the Exome Aggregation Consortium (ExAC) database [20] and 76,314 European-ancestry individuals from a genetic testing laboratory (Counsyl) (see Methods). Dashed blue line indicates cases where allele frequency in Counsyl is the same as in ExAC.

**Fig S2.**
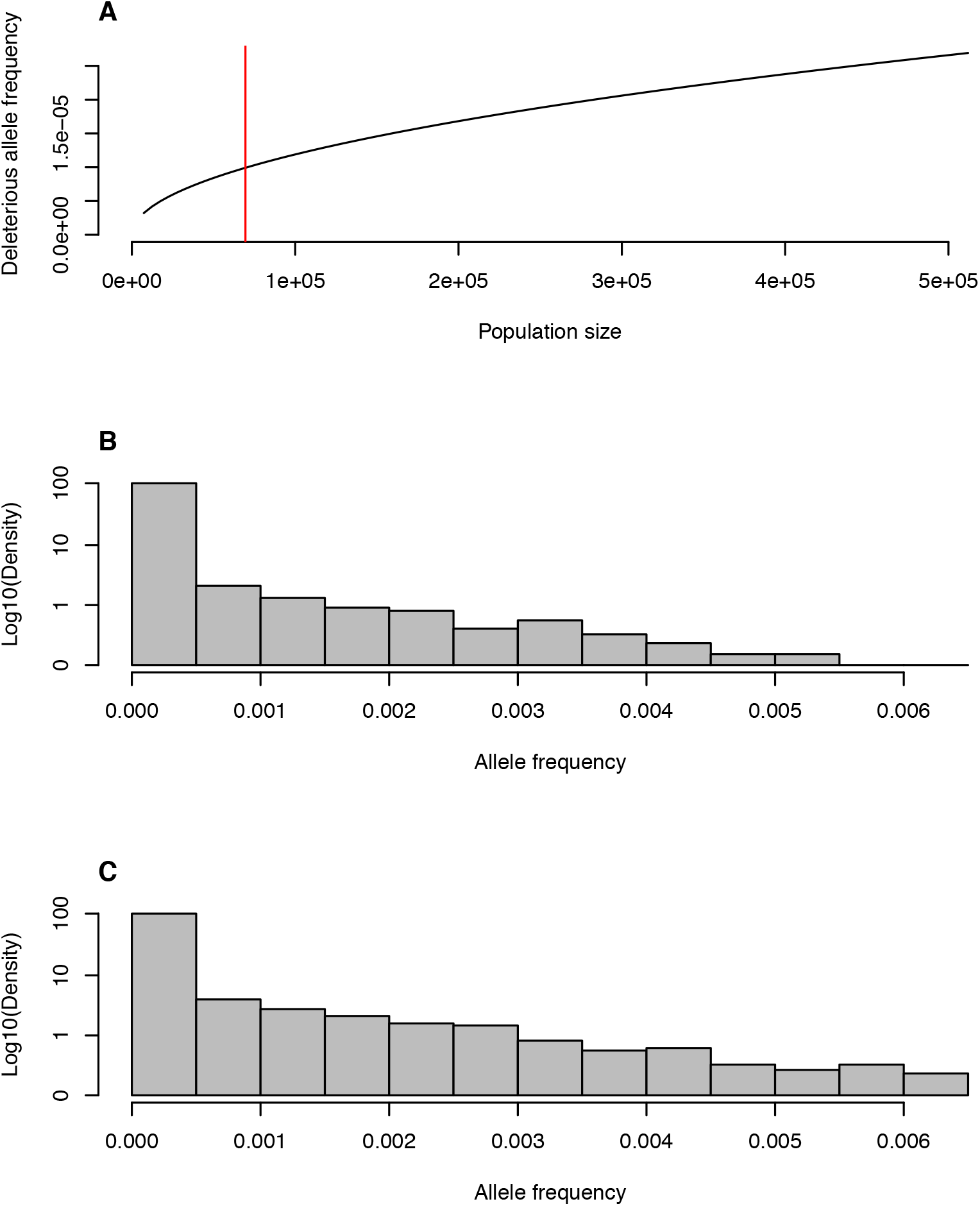
Comparisons of mutation-selection balance models for finite population sizes. (A) Mean allele frequency as a function of effective population size, under a model of constant population size. The X-axis range corresponds to the minimum and maximum effective population size estimated in [22]. The red bar indicates the value of a constant population size at which the mean allele frequency predicted is the same as the mean allele frequency estimated in simulations, for an average mutation rate of 1.5 ×10^−8^ per bp per generation [31]. (B-C) The allele frequency distribution (in grey) is presented for 100,000 observations based on (B) the complex demographic scenario inferred by Tennessen et al. [22] for the evolution of European populations based on simulations (see Methods) and on (C) the finite, constant size population model, with *N* set to 69,537 individuals to match the mean allele frequency with (B). Both models assume complete lethality (*s*=1) and recessivity (*h* = 0).

**Fig S3.**
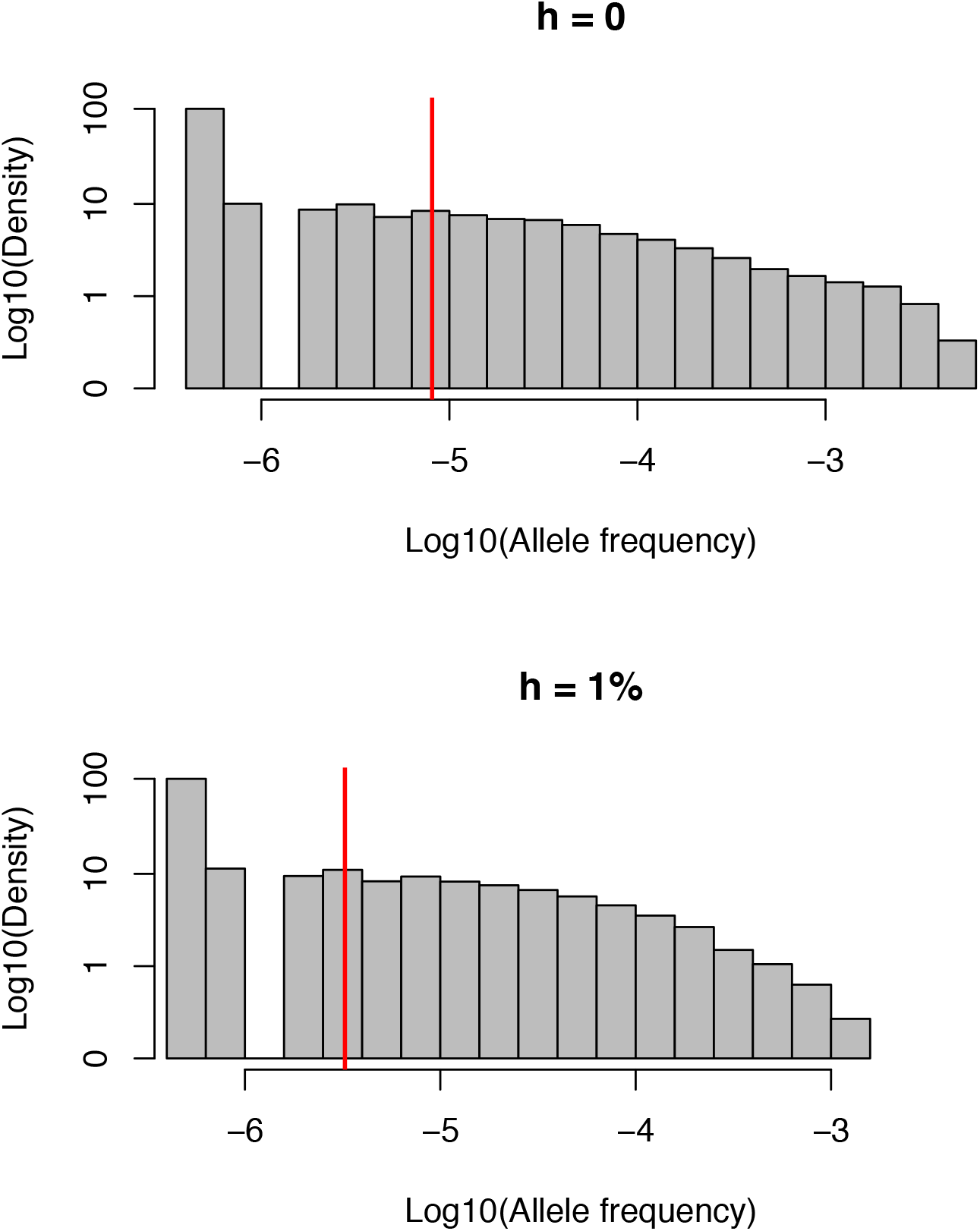
The impact on disease allele frequencies of a small fitness effect in heterozygotes (*h*=0.01). Shown is the distribution of deleterious allele frequencies generated from 100,000 simulations in each case. Means are represented by red vertical bars. For visualization, an allele frequency *q*=0 is set to 0.5 × 10^−6^. When a small fitness effect in heterozygotes is considered in the simulations, the mean allele frequency decreases by 61% relative to no effect. The two distributions differ significantly by a Kolmogorov-Smirnov test (p-value < 10^−15^). The mutation rate *u* was set to 1.5 × 10^−8^ per bp per generation, reflective of the mean mutation rate estimated for exons [31].

**Fig S4.**
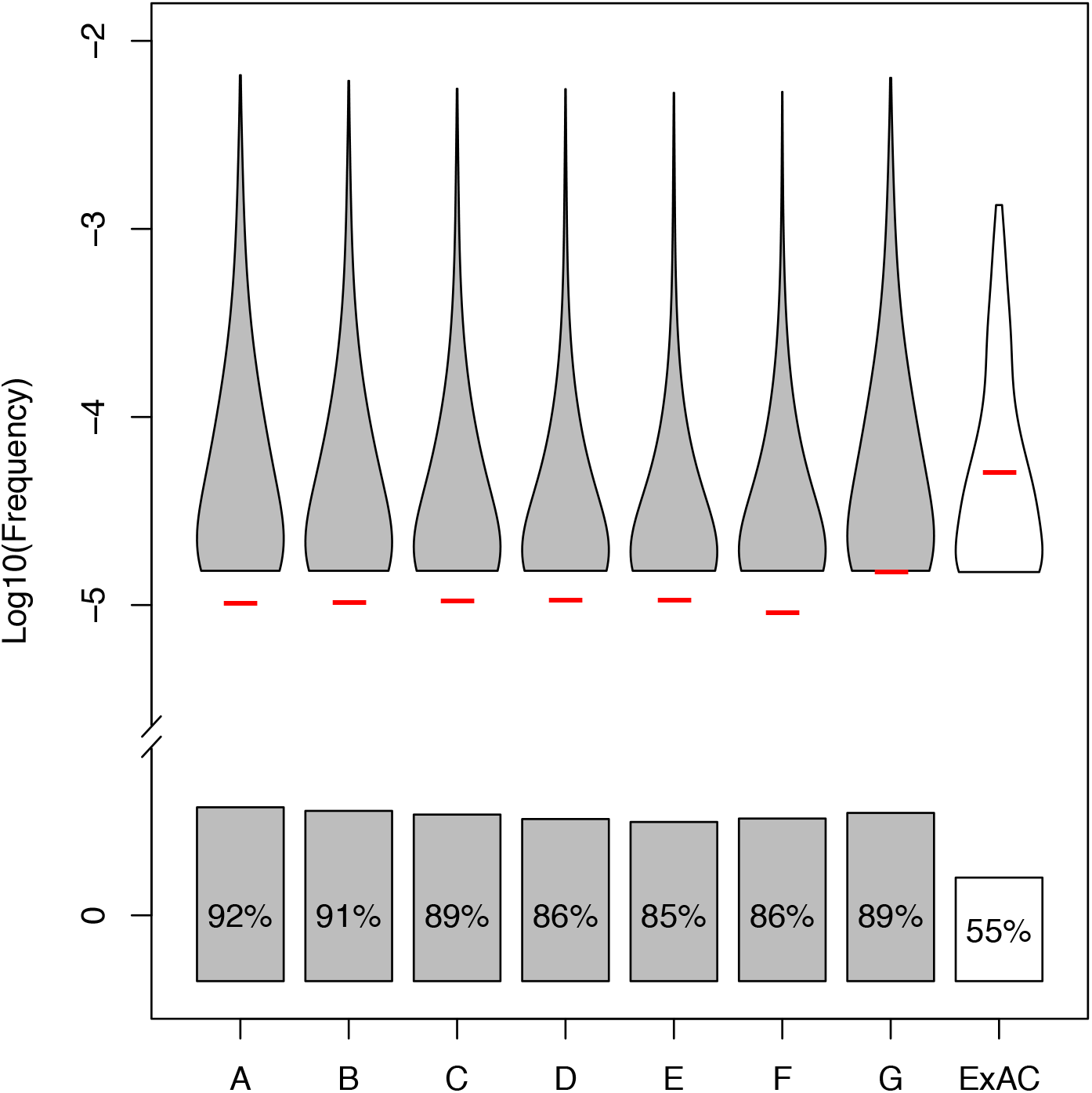
Effect of varying the end population size and the average mutation rate on the sample frequency of recessive, lethal mutations. Tennessen et al. [22] inferred the present effective population size of Europeans to be 512,000 individuals (column A). We considered the effect of larger present population sizes (2-, 4-, 10-, and 20-fold increase, denoted by columns B, C, D and E respectively), keeping other demographic parameters the same. We also included a model (F) where rapid growth begins immediately after the out-of-Africa bottleneck, representing a more extreme scenario of population growth in comparison to the two-stage and more gradual scenario proposed by Tennessen et al. (2012). For A-F we use the average *u* = 1.5 × 10^−8^ per bp per generation [31]. Model G considers a larger *u* (2.25 × 10^−8^, i.e., a 1.5-fold increase from A-F), with all other parameters (e.g., variance in mutation rates across simulations, the demographic model) kept the same as in column A. The observed allele frequency distribution of 385 disease mutations in ExAC is shown in white. Violin plots show the density distribution of the log_10_ of the frequency of alleles segregating in these samples, whereas boxes indicate the proportion of sites for which the deleterious mutation was not observed. All distributions differ significantly from one another (i.e., all p-values are < 10^−15^ by a Kolmogorov-Smirnov test).

**Fig S5.**
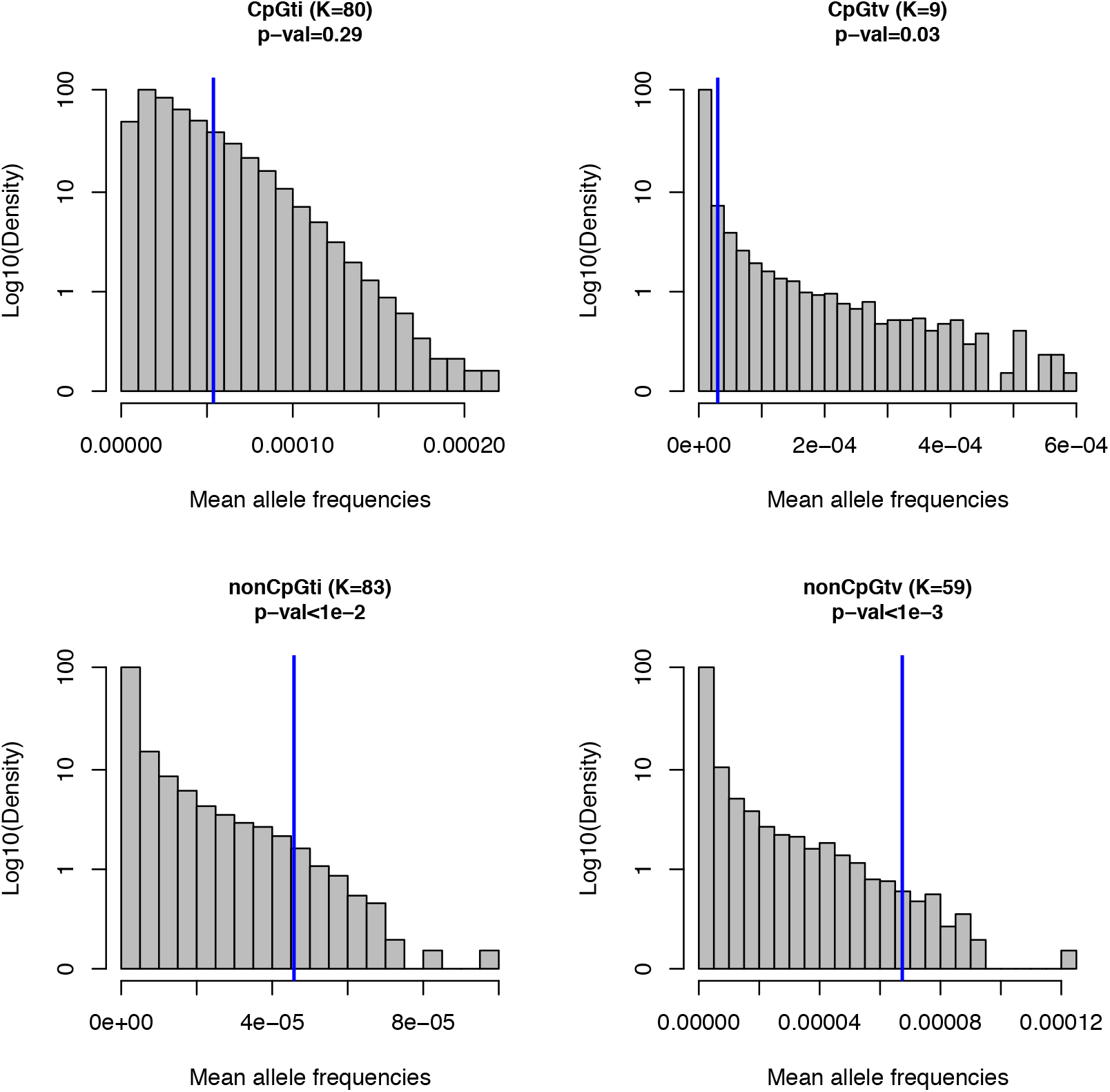
Expected distribution and the observed mean allele frequencies of recessive, lethal disease mutations (excluding mutations in *CFTR* and *DHCR7*). As in Fig 2, the four panels correspond to four different mutation types. The title of the panel indicates the mutation type, followed by *N*, the total number of mutations of that type, with p-values for the difference between observed and expected mean frequencies below. Distributions in grey are for 100,000 observations of the expected mean allele frequencies across *K* mutations, and were obtained from simulations based on a plausible demographic model for European populations [22] (see Methods). Blue bars represent the observed values estimated from 33,370 individuals of European ancestry from ExAC. As opposed to in Fig 2, here, we did not include mutations present in two genes (*CFTR* and *DHCR7*) that were outliers in the gene-level analysis (Fig 3) and were reported elsewhere to be carried by healthy homozygous individuals [21].

**Fig S6.**
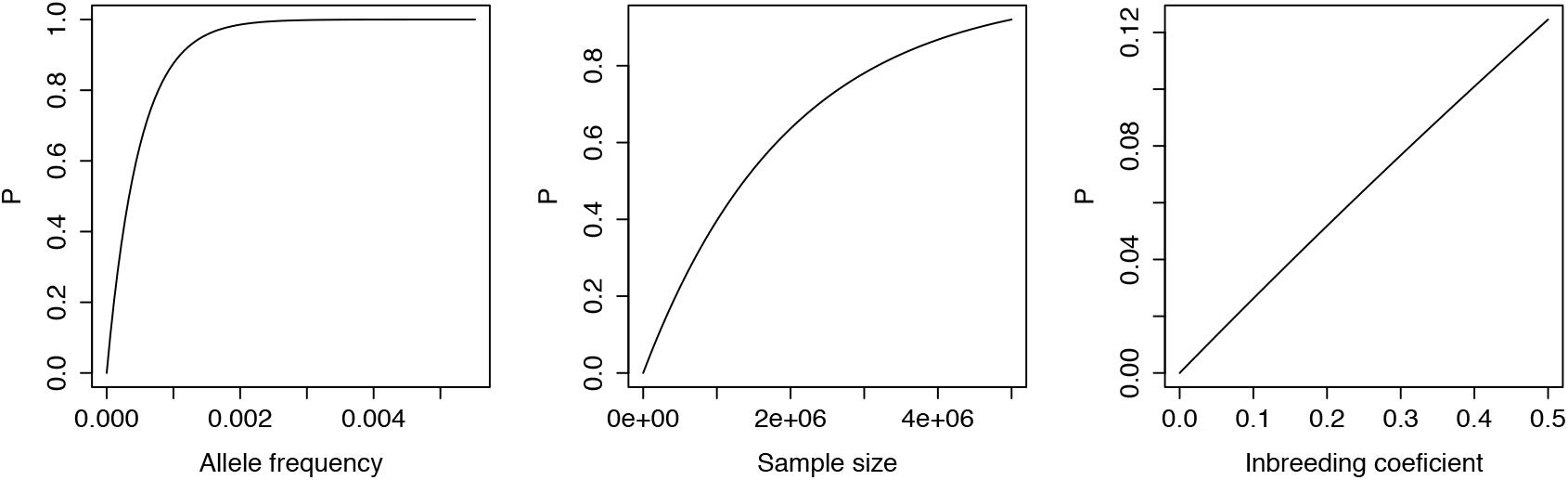
The probability of a mutation being ascertained, given its allele frequency *p*, the sample size *n_a_* of the putative ascertainment study and the inbreeding coefficient *F_a_* in the population in which the ascertainment study was conducted. In each case, we let only one parameter (*p, n_a_* or *F_a_*) to vary, while fixing the others at *p*=1×10^−5^ (corresponding to the mean allele frequency from simulations), *n_a_*=10,000, and *F_a_*=1/16 (corresponding to marriage between first cousins, a plausible scenario for a population with widespread inbreeding).

**Fig S7.**
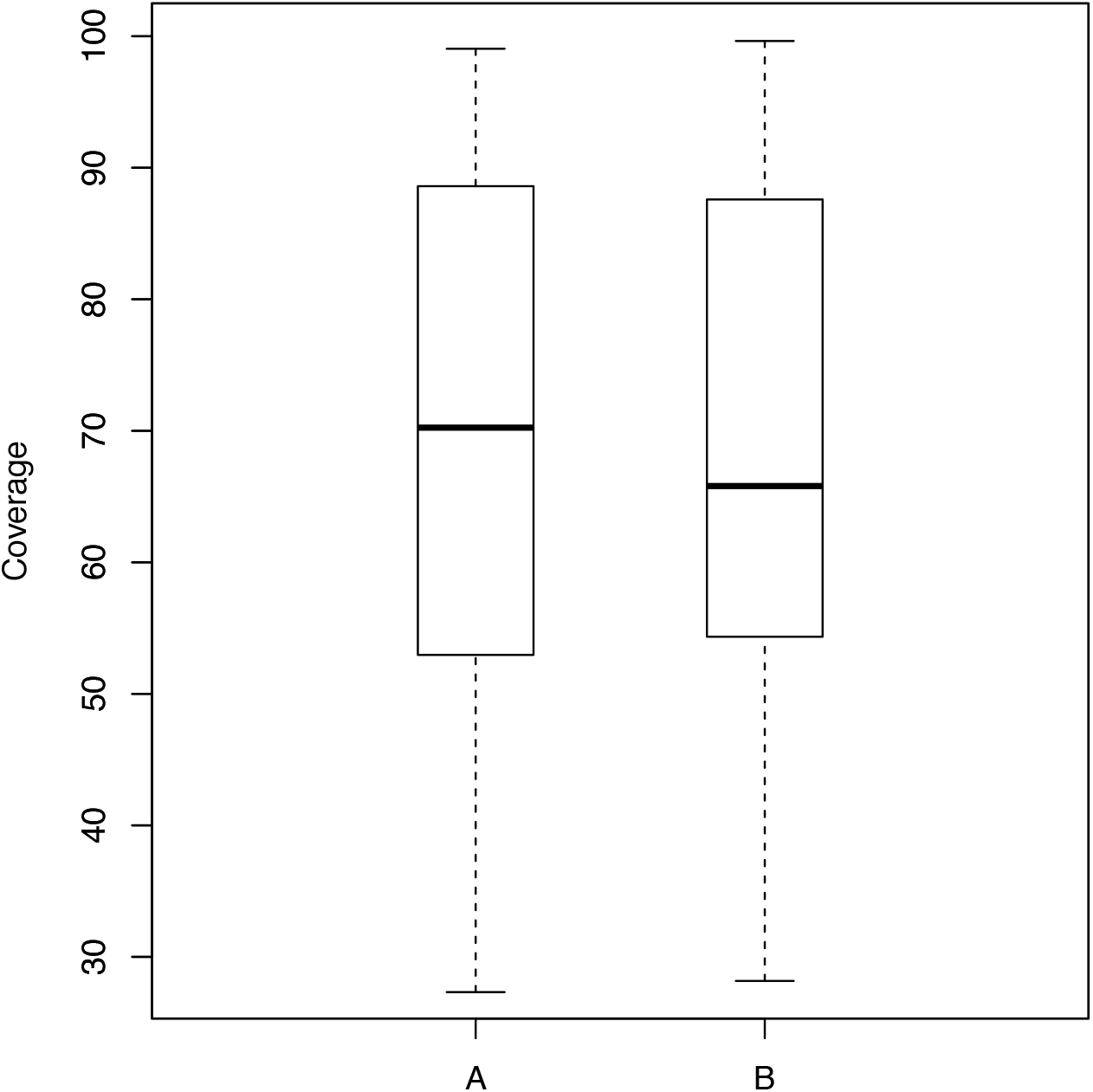
Depth of coverage for 385 mutations in ExAC known to cause lethal, Mendelian diseases. Box plots show the mean (black bar) and the lower and upper quartiles for (A) the 248 sites with non-zero sample frequencies in ExAC, for which the number of sequenced non-Finnish European individuals was reported (*n* = 32,881) and (B) the 137 sites for which we did not have this information. Since distributions of depth of coverage are similar between the two sets, we assumed that 32,881 individuals were sequenced at all sites, and used this number to subsample simulations to match the sample size of the ExAC data.

## References

1. Simons YB, Turchin MC, Pritchard JK, Sella G (2014) The deleterious mutation load is insensitive to recent population history. Nat Genet 46: 220–224.

2. Simons YB, Sella G (2016) The impact of recent population history on the deleterious mutation load in humans and close evolutionary relatives. Curr Opin Genet Dev 41: 150–158.

3. Nei M (1968) The frequency distribution of lethal chromosomes in finite populations. Proc Natl Acad Sci U S A 60: 517–524.

4. Gillespie JH (2004) Population Genetics: A Concise Guide. Baltimore, MD: Johns Hopkins University Press.

5. Brandvain Y, Wright SI (2016) The Limits of Natural Selection in a Nonequilibrium World. Trends Genet 32: 201–210.

6. Balick DJ, Do R, Cassa CA, Reich D, Sunyaev SR (2015) Dominance of Deleterious Alleles Controls the Response to a Population Bottleneck. PLoS Genet 11: e1005436.

7. Beauchamp KA, Muzzey D, Wong KK, Hogan GJ, Karimi K, et al. (2016) Systematic Design and Comparison of Expanded Carrier Screening Panels. bioRxiv.

8. Cormet-Boyaka E, Jablonsky M, Naren AP, Jackson PL, Muccio DD, et al. (2004) Rescuing cystic fibrosis transmembrane conductance regulator (CFTR)-processing mutants by transcomplementation. Proc Natl Acad Sci U S A 101: 8221–8226.

9. Rapino D, Sabirzhanova I, Lopes-Pacheco M, Grover R, Guggino WB, et al. (2015) Rescue of NBD2 mutants N1303K and S1235R of CFTR by small-molecule correctors and transcomplementation. PLoS One 10: e0119796.

10. Andressoo JO, Jans J, de Wit J, Coin F, Hoogstraten D, et al. (2006) Rescue of progeria in trichothiodystrophy by homozygous lethal Xpd alleles. PLoS Biol 4: e322.

11. Gallati S (2014) Disease-modifying genes and monogenic disorders: experience in cystic fibrosis. Appl Clin Genet 7: 133–146.

12. Corvol H, Blackman SM, Boelle PY, Gallins PJ, Pace RG, et al. (2015) Genome-wide association meta-analysis identifies five modifier loci of lung disease severity in cystic fibrosis. Nat Commun 6: 8382.

13. Habara A, Steinberg MH (2016) Minireview: Genetic basis of heterogeneity and severity in sickle cell disease. Exp Biol Med (Maywood) 241: 689–696.

14. Hedrick PW (2011) Population genetics of malaria resistance in humans. Heredity (Edinb) 107: 283–304.

15. Chong JX, Buckingham KJ, Jhangiani SN, Boehm C, Sobreira N, et al. (2015) The Genetic Basis of Mendelian Phenotypes: Discoveries, Challenges, and Opportunities. Am J Hum Genet 97: 199–215.

16. Kong A, Frigge ML, Masson G, Besenbacher S, Sulem P, et al. (2012) Rate of de novo mutations and the importance of father's age to disease risk. Nature 488: 471–475.

17. Turner TN, Douville C, Kim D, Stenson PD, Cooper DN, et al. (2015) Proteins linked to autosomal dominant and autosomal recessive disorders harbor characteristic rare missense mutation distribution patterns. Hum Mol Genet 24: 5995–6002.

18. Cooper DN, Youssoufian H (1988) The CpG dinucleotide and human genetic disease. Hum Genet 78: 151–155.

19. Akalin N, Zietkiewicz E, Makalowski W, Labuda D (1994) Are CpG sites mutation hot spots in the dystrophin gene? Hum Mol Genet 3.

20. Lek M, Karczewski KJ, Minikel EV, Samocha KE, Banks E, et al. (2016) Analysis of protein-coding genetic variation in 60,706 humans. Nature 536: 285–291.

21. Chen R, Shi L, Hakenberg J, Naughton B, Sklar P, et al. (2016) Analysis of 589,306 genomes identifies individuals resilient to severe Mendelian childhood diseases. Nat Biotechnol 34: 531–538.

22. Tennessen JA, Bigham AW, O'Connor TD, Fu W, Kenny EE, et al. (2012) Evolution and functional impact of rare coding variation from deep sequencing of human exomes. Science 337: 64–69.

23. Wright S (1937) The Distribution of Gene Frequencies in Populations. Proc Natl Acad Sci U S A 23: 307–320.

24. Harpak A, Bhaskar A, Pritchard JK (2016) Mutation Rate Variation is a Primary Determinant of the Distribution of Allele Frequencies in Humans. PLoS Genet 12: e1006489.

25. Browning Sharon R, Browning Brian L (2015) Accurate Non-parametric Estimation of Recent Effective Population Size from Segments of Identity by Descent. The American Journal of Human Genetics 97: 404–418.

26. Reich DE, Lander ES (2001) On the allelic spectrum of human disease. Trends Genet 17: 502–510.

27. Pritchard JK, Cox NJ (2002) The allelic architecture of human disease genes: common disease-common variant…or not? Hum Mol Genet 11: 2417–2423.

28. Zwick ME, Cutler DJ, Chakravarti A (2000) Patterns of genetic variation in Mendelian and complex traits. Annu Rev Genomics Hum Genet 1: 387–407.

29. Cassa CA, Weghorn D, Balick DJ, Jordan DM, Nusinow D, et al. (2017) Estimating the Selective Effect of Heterozygous Protein Truncating Variants from Human Exome Data. Nat Genet 49: 806–810.

30. The 1000 Genomes Consortium Project (2015) A global reference for human genetic variation. Nature 526: 68–74.

31. Neale BM, Kou Y, Liu L, Ma/'ayan A, Samocha KE, et al. (2012) Patterns and rates of exonic de novo mutations in autism spectrum disorders. Nature 485: 242–245.

32. Fu W, O'Connor T, Jun G, Kang H, Abecasis G, et al. (2013) Analysis of 6,515 exomes reveals the recent origin of most human protein-coding variants. Nature 493: 216–220.

33. Lohmueller KE, Indap AR, Schmidt S, Boyko AR, Hernandez RD, et al. (2008) Proportionally more deleterious genetic variation in European than in African populations. Nature 451: 994–997.

34. Peischl S, Dupanloup I, Bosshard L, Excoffier L (2016) Genetic surfing in human populations: from genes to genomes. Curr Opin Genet Dev 41: 53–61.

35. Pritchard JK, Cox NJ (2002) The allelic architecture of human disease genes: common disease-common variant…or not? Hum Mol Genet 11:2417–23.

36. Segurel L, Wyman MJ, Przeworski M (2014) Determinants of mutation rate variation in the human germline. Annu Rev Genomics Hum Genet 15: 47–70.

37. Aggarwala V, Voight BF (2016) An expanded sequence context model broadly explains variability in polymorphism levels across the human genome. Nat Genet 48: 349–355.

38. Hodgkinson A, Eyre-Walker A Variation in the mutation rate across mammalian genomes. Nat Rev Genet 12: 756–766.

39. Moorjani P, Amorim CE, Arndt PF, Przeworski M (2016) Variation in the molecular clock of primates. Proc Natl Acad Sci U S A 113: 10607–10612.

40. Kamphans T, Sabri P, Zhu N, Heinrich V, Mundlos S, et al. (2013) Filtering for compound heterozygous sequence variants in non-consanguineous pedigrees. PLoS One 8: e70151.

41. Xue Y, Chen Y, Ayub Q, Huang N, Ball EV, et al. (2012) Deleterious- and disease-allele prevalence in healthy individuals: insights from current predictions, mutation databases, and population-scale resequencing. Am J Hum Genet 91: 1022–1032.

42. Piton A, Redin C, Mandel JL (2013) XLID-causing mutations and associated genes challenged in light of data from large-scale human exome sequencing. Am J Hum Genet 93: 368–383.

43. Gazave E, Ma L, Chang D, Coventry A, Gao F, et al. (2014) Neutral genomic regions refine models of recent rapid human population growth. Proc Natl Acad Sci U S A 111: 757–762.

44. Gao F, Keinan A (2016) Explosive genetic evidence for explosive human population growth. Curr Opin Genet Dev 41: 130–139.

45. Wright S (1931) Evolution in Mendelian populations. Genetics 16: 97–159.

46. Ober C, Hyslop T, Hauck WW (1999) Inbreeding Effects on Fertility in Humans: Evidence for Reproductive Compensation. The American Journal of Human Genetics 64: 225–231.

47. Gabriel SE, Brigman KN, Koller BH, Boucher RC, Stutts MJ (1994) Cystic fibrosis heterozygote resistance to cholera toxin in the cystic fibrosis mouse model. Science 266: 107–109.

48. Wagener D, Cavalli-Sforza LL, Barakat R (1978) Ethnic variation of genetic disease: roles of drift for recessive lethal genes. American Journal of Human Genetics 30: 262–270.

49. Quinton PM (1994) Human genetics. What is good about cystic fibrosis? Curr Biol 4: 742–743.

50. Chakravarti A, Chakraborty R (1978) Elevated frequency of Tay-Sachs disease among Ashkenazic Jews unlikely by genetic drift alone. American Journal of Human Genetics 30: 256–261.

51. Ewens WJ (1978) Tay-Sachs disease and theoretical population genetics. American Journal of Human Genetics 30: 328–329.

52. Richards S, Aziz N, Bale S, Bick D, Das S, et al. (2015) Standards and guidelines for the interpretation of sequence variants: a joint consensus recommendation of the American College of Medical Genetics and Genomics and the Association for Molecular Pathology. Genet Med 17: 405–423.

53. Landrum MJ, Lee JM, Riley GR, Jang W, Rubinstein WS, et al. (2014) ClinVar: public archive of relationships among sequence variation and human phenotype. Nucleic acids research 42: D980–D985.

54. Eppig JT, Blake JA, Bult CJ, Kadin JA, Richardson JE (2015) The Mouse Genome Database (MGD): facilitating mouse as a model for human biology and disease. Nucleic Acids Res 43: D726–736.

55. Haque IS, Lazarin GA, Kang H, Evans EA, Goldberg JD, et al. (2016) MOdeled fetal risk of genetic diseases identified by expanded carrier screening. JAMA 316: 734–742.

56. Mugal CF, Ellegren H (2011) Substitution rate variation at human CpG sites correlates with non-CpG divergence, methylation level and GC content. Genome Biol 12: R58.

57. North BV, Curtis D, Sham PC (2002) A Note on the Calculation of Empirical Values from Monte Carlo Procedures. The American Journal of Human Genetics 71: 439–441.

58. Quinlan AR, Hall IM (2010) BEDTools: a flexible suite of utilities for comparing genomic features. Bioinformatics 26: 841–842.

59. R Core Team (2015) R: A Language and Environment for Statistical Computing. Vienna, Austria.

60. Online Mendelian Inheritance in Man O (2016). Baltimore, MD: McKusick-Nathans Institute of Genetic Medicine, Johns Hopkins University.

